# The MYC axis in advanced prostate cancer is impacted through concurrent targeting of ERβ and AR using a novel ERβ-selective ligand alongside Enzalutamide

**DOI:** 10.1101/2023.11.15.567282

**Authors:** Jaimie S. Gray, Sajad A. Wani, Shahid Hussain, Phoebe Huang, Debasis Nayak, Mark D. Long, Clayton Yates, Steven K. Clinton, Chad E. Bennet, Christopher C. Coss, Moray J. Campbell

## Abstract

We have dissected the role of Estrogen receptor beta (ERβ) in prostate cancer (PCa) with a novel ERβ ligand, OSU-ERb-12. Drug screens revealed additive interactions between OSU-ERB-12 and either epigenetic inhibitors or the androgen receptor antagonist, Enzalutamide (Enza). Clonogenic and cell biolody studies supported the potent additive effects of OSU-ERB-12 (100nM) and Enza (1µM). The cooperative behavior was in PCa cell lines treated with either OSU-ERB-12 plus Enza or combinations involving 17β-estradiol (E2). OSU-ERb-12 plus Enza uniquely impacted the transcriptiome, accessible chromatin, and the AR, MYC and H3K27ac cistromes. This included skewed transcriptional responses including suppression of the androgen and MYC transcriptomes, and repressed MYC protein. OSU-ERb-12 plus Enza uniquely impacted chromatin accessibility at approximately 3000 nucleosome-free sites, enriched at enhancers, enriched for basic Helix-Loop-Helix motifs. CUT&RUN experiments revealed combination treatment targeting of MYC, AR, and H3K27ac again shaping enhancer accessibility. Specifically, it repressed MYC interactions at enhancer regions enriched for bHLH motifs, and overlapped with publicly-available bHLH cistromes. Finally, cistrome-transcriptome analyses identified ∼200 genes that distinguished advanced PCa tumors in the SU2C cohort with high androgen and low neuroendocrine scores.

**Statement of Implication:** Targeting ERβ has potentially to augment AR antagonism to restrain MYC signaling and limit growth of advanced prostate cancer.

## INTRODUCTION

Prostate Cancer (PCa) patients can experience treatment failure following surgical or radiotherapy treatments their disease often progresses. In this PCa stage androgen receptor (AR) signaling is targeted with androgen deprivation therapy (ADT). Unfortunately, the impact of ADT is often short-lived, and the disease progresses towards ADT resistant PCa (ADT-R-PCa), associated with the neuroendocrine (NE) and other androgen-indifferent states, which are frequently lethal. One mechanism of resistance to ADT is the interplay of dominant transcription factors such as MYC(1) and ONECUT2(2) that combine with altered epigenomic states to allow transition to alternative lineages (reviewed in(3)). Therefore, there is a need to understand mechanisms that promote normal AR functions to promote differentiation, and which may sustain the effectiveness of ADT. For example, ADT has been examined in combination with prednisone, a glucocorticoid receptor ligand(4).

One alternative approach is to consider other nuclear receptors that interact with the AR, such as the estrogen receptor beta (ERβ). The therapeutic benefit of estrogens to treat PCa was established in the 1960s(5) but stopped due to cardiac related side effects. Perhaps hinting at a functional role for estrogens in the prostate, the ERβ was cloned from the rat prostate(6). ERβ knockout mice display increased prostatic hyperplasia, fibrosis or inflammation in both ventral prostate and mammary glands(7,8). ERβ also exerts anti-proliferative and pro-apoptotic actions in the central nervous system, colon, and prostate(9). Subsequent analyses has supported ERβ functioning as a growth suppressor in breast and ovarian cancers(10), and higher expression of *ESR2* is correlated with better overall survival outcomes in both prostate and breast cancers(11). There is also evidence to suggest that exon skipping can generate ERβ variants with a dominant negative effect(12). Collectively, these findings support the possibility of targeting estrogenic signaling in PCa.

There is a complex interplay between ERβ and ERα, which form both homodimers and heterodimers, and both participate in cytoplasmic signal transduction events. In the nucleus they function both in *cis* and *trans* to regulate transcription, and to regulate splicing events(13). Studying this complex biology is compounded by availability of ERβ selective ligands, although recently a number have been created with differing potency and selectivity of ERβ over Erα, including LY3201 (Diarylproprionitrile, DPN) and LY500307 (Erteberel). LY3201 treatment downregulates AR and target genes in PCa cell lines(14) and DPN treatment maintains an epithelial phenotype(15). We have developed novel para-carborane ERβ-selective agonists with much improved pharmacodynamics compared to these other compounds and one, OSU-ERb-12, has over 200-fold functional selectivity for ERβ over ERα(16).

We have examined the functions of targeting ERβ via OSU-ERb-12 in combination with other drugs known to potentiate nuclear receptor signaling, including ADT drugs. Given that nuclear receptors interact with a large number of coregulator proteins to control chromatin states, we also examined how epigenetic drugs such as HDAC inhibitors can potentiate via OSU-ERb- 12 (17-20). This approach identified that OSU-ERb-12 and the ADT drug Enzalutamide (Enza) display potent additive interactions in cell biology experiments. Subsequently, integrative genomic approaches revealed a mechanism by which OSU-ERb-12 and Enza lead to repression of chromatin accessibility at enhancer sites for MYC and repressed MYC signaling.

## MATERIALS AND METHODS

### Cell culture and materials

Cells utilized were the non-malignant prostate cells HPr1AR (gift from Eric Bolton, University of Illinois, Chicago, IL, USA) and malignant LNCaP, 22Rv1, LNCaP C4-2, PC-3 and DU145 were obtained from the ATCC. All cells were maintained at 37°C and 5.0% CO_2_ using a cell culture incubator with UV contamination control. HPr1AR cells were maintained in keratinocyte serum free media (Gibco) supplemented with 25mg of Bovine pituitary extract (BPE), 10ug human epithelial growth factor (hEGF) and 10% FBS (Sigma); LNCaP, LNCaP C4-2, 22Rv1, PC-3 and DU145 cells in RPMI 1640 Medium (Gibco) supplemented with 10% FBS (Sigma). Non-malignant African American prostate cells, RC43N and isogenic PCa RC43T(21,22) and isogenic AA PCa (RC43T, RC77T) were maintained in keratinocyte serum free media (supplemented with 25mg of Bovine pituitary extract, 10ug EGF and 10% FBS). For experimental media cells were cultured in phenol red free conditions. Cell lines were authenticated by STR profiling and confirmed mycoplasma free by RT-PCR routinely.

### CRISPR ERβ knockout

crRNAs oligos (Integrated DNA Technologies (IDT) designer tool) targeted to *ESR1* and *ESR2* were added, tracrRNA (*IDT*) and nuclease buffer and incubated with Cas9, and Cas9 plus reagent (IDT) with OPtiMEM media (Gibco) to form the ribonucleic protein complex (RNP), which combined with CRISPRMAX lipofectamine reagent (Invitrogen) to form the lipo-RNP complex and added to 22Rv1 cells and incubated overnight. After clonal isolation knockout confirmation was confirmed by Sanger sequencing to generate the ERα knockout (ERKO) and ERβ knockout (BERKO) cells.

### Doubling time assay

Parental, ERKO and BERKO 22Rv1 cells were seeded (0.1 x 10^6^ cells/well) and viable cells counted every 24 h and analyzed using doubling time formulas.

### Scratch Assay

Parental, ERKO and BERKO 22Rv1 cells were seeded were seeded (0.4 x 10^6^ cells/well) and after 24 h the wells were scratched with a sterile pipet. At indicated time points images were captured and area of the scratch calculated using ImageJ software.

### OSU-ERb-12 combination drug screens

OSU-ERb-12 was generated in the Ohio State University Drug Development Institute, and all other drugs were commercially available. Diluent, stock concentration and source are listed in **Supplementary Table 1**. Multi-dose screens were undertaken in LNCaP, LNCaP C4-2, 22Rv1, PC-3, and DU145 by plating mid-exponentially proliferating cells (1750 cells per well/384 well plate) in phenol red free media 24 h prior to addition of drugs through an automated robotic liquid handler (BioMek, Beckman Coulter). After 72h ATP-luciferase assay reagent was added (CellTiter-Glo, Promega) to measure ATP activity per well (Synergy H10, BioTek).

### Drug screen analysis

Raw light units were normalized to negative controls on each plate and percent viability determined. Contour plots were created using `lattice`, in R. Concentrations of each drug in the combination were plotted on each axis; x for OSU-ERb-12, y for drug library compound, and z axis represents viability, which is plotted using a color scale, with lighter color representing inhibition of viability. Delta values were calculated by taking the additive value of each single agent’s inhibition (expected value) subtracted from the observed inhibition value of the combination. Positive delta values indicate increased inhibition above additive assumption vs negative delta values indicate a drug antagonism and the p values calculated by comparing the expected versus the observed values, as previously(19).

### Clonogenic assays

Colony formation was measured in the presence of OSU-ERb-12 (100nM), Enza (1μM), EGCG (10 μM), or FK228 (1nM), or combinations with OSU-ERb-12 by plating 100 cells in each well of a 6 well plate and treated with single agents or combinations every three days for a period of 14 days. After 14 days cells were washed and fixed with neutral buffered formalin and stained with crystal violet stain and quantified as previously reported(23)

### Cell cycle and apoptosis assays

Cells were grown in the presence of OSU-ERb-12 (100nM), Enza (1uM), or OSU-ERb-12 (100nM) + Enza (1uM) for 48h and processed as previously(24) reported using Annexin-V-FLUOS staining kit (Roche) and Propidium Iodide, respectively.

### RT-qPCR

Total RNA was isolated from exponentially growing cells via E.Z.N.A total RNA kit I (Omega BioTek) and converted to cDNA with high-capacity cDNA Reverse Trancriptase kit (Applied Biosciences), following manufacturer’s protocols and gene expression quantified via Applied Biosystems Quant Studio 6/7 Real-Time PCR System (Applied Biosystems), for TaqMan (Thermo Fisher Scientific) applications. All targets were detected using pre-designed TaqMan Gene Expression Assays (Thermo Fisher Scientific); *ESR1, ESR2, AR, TMPRSS2, TP53, MYC, FOXA1*. All RT-qPCR experiments were performed in biological triplicates, with at least technical triplicates. Fold changes were determined using the 2-ΔΔCt method as the difference between experimental group and respective control group.

### Western Immunoblotting

Total cellular protein was harvested from exponentially growing cells, treated as indicated, washed in ice cold PBS. Cell lysis was in ice cold RIPA buffer (50mM Tris-HCl pH 7.4, 150mM NaCl, 1% v/v Triton X-100, 1mM EDTA pH 8.0, 0.5% w/v sodium deoxychlorate, 0.1% w/v SDS) containing 1x cOmplete Mini Protease Inhibitor Tablets (Roche).

Protein concentrations were quantified using DC Protein Assay (Bio-Rad). Equal amounts of proteins (30-60µg) were resolved via SDS polyacrylamide gel electrophoresis (SDS-PAGE) using precast polyacrylamide gradient gels (Mini-Protean TGX, Bio-Rad) and transferred onto polyvinylidene fluoride (PVDF) membrane (Roche) for 30V for 16h. Post transfer, membranes were blocked with 5% non-fat dry milk (NFDM) for 1 hour at room temperature. Blocked membranes were probed with primary antibody against MYC (18583, Cell Signaling Technology), AR (5153, Cell Signaling Technology), ARv7 (198394, abcam), ERα (8644, Cell Signaling Technology), ERβ (89954S, Cell Signaling Technology) Flag tag (14793, Cell Signaling Technology; F1804, Sigma), Cytokeratin 5 (AB_2538529, Thermofisher), Cytokeratin 8 (ab53280, abcam) overnight at 4C. Primary antibody was detected after probing for 1h with HRP-linked rabbit anti-mouse IgG (AB_228307, Thermofisher) or goat anti-rabbit IgG (AB_228341, Thermofisher) secondary antibody at room temperature using Chemiluminescent Western Blotting substrate (Pierce). Signal quantification was performed using the ProteinSimple Fluorochem M Imager(24).

### RNA-Seq

LNCaP, LNCaP C4-2 and 22Rv1 cells (0.5×10^6^) were plated overnight and subsequently exposed to the indicated treatments for 12h and 48h in biological triplicate samples and analyzed by RNA-Seq, of 1ug total RNA prepared into libraries prepared with the TruSeq Stranded Total RNA kit (Illumina Inc). Alignment of raw sequence reads to the human transcriptome (hg38) was performed via `Rsubread` (25) and transcript abundance estimates were normalized and differentially expressed genes (DEGs) identified using a standard `edgeR` pipeline(26). Functional annotation of gene sets: Pathway enrichment analysis and gene set enrichment analysis (`GSEA`) were performed using gene sets from the Molecular signatures database (MSigDB). For transcript-aware analyses, the FASTQ files were aligned with `salmon`(27) and differentially enriched transcripts were identified using `DRIMSeq`(28) in a similar workflow to `edgeR`.

### ATAC-Seq

22Rv1 cells (5×10^4^) were plated overnight and subsequently exposed to the indicated treatments for 6h, after which cells were washed twice with PBS and collected by trypsinization. Pelleted cells were resuspended in 50ul of ATAC-resuspension buffer (ATAC-RSB - 10mM Tris-HCl, 10mM NaCl, 3mM MgCl_2_) containing (0.1% NP-40, 0.1% tween-20, and 0.01% digitonin) and pipetted up and down 3 times. Further, 1ml of ATAC-wash-resuspension buffer (ATAC-RSB + 0.1% tween 20) was used to pellet down the nuclei. The nucleic were further resuspended in transposition mix (Tagment DNA buffer (Illumina), PBS, Digitonin 0.01%, tween 20 0.1%, nuclease free water, and transposase 2.5ul/reaction (Illumina). After transposase reaction, samples were cleaned up using Monarch PCR and DNA cleanup kit (New England Biosciences). Samples were then amplified in a PCR reaction using NEBNext PCR Master Mix (New England Biosciences) and Nextera compatible indexing primers, Unique Dual Index configuration, (Integrated DNA Technologies). After amplification, samples were cleaned up using AMPure beads (Beckman Coulter) to isolate fragments between 100bp and 1000bp, and sequenced (29). Alignment of reads to the human transcriptome (hg38) was performed via `Rsubread` (25) and the ATAC-Seq data separated into nucleosome free (NF), mono-, di-and tri-nucleosome compartments (`ATACSeqQC`)(30), and differential enrichment measured by `csaw`(31).

### CUT&RUN

22Rv1 cells (0.5×10^6^) were plated overnight and subsequently exposed to the indicated treatments for 6h. CUT&RUN was performed. Briefly, approximately cells were adsorbed to concanavalin A beads (Epicypher), and washed (150mM NaCl, 20mM HEPES, 0.01% Digitonin, 0.5mM Spermidine, protease inhibitor cocktail EDTA-free mini tab 1tab/10mL (Roche)) to permeabilize cells and nuclei. Samples were incubated overnight with antibodies against IgG (2729, Cell Signaling Technology), H3K27 (8173, Cell Signaling Technology), AR (5153, Cell Signaling Technology), ARv7 (ab198394, abcam), MYC (9402, Cell Signaling Technology) for 16 hours, and antibody bound DNA was then enzymatically cut with pAG-MNase (Epicypher). Samples were treated with Stop Buffer (340mM NaCl, 20mM EDTA, 4mM EGTA, 50ug/mL RNase A, 50ug/mL Glycogen) to prevent further enzymatic activity. DNA fragments released into supernatant were collected and purified by Monarch PCR and DNA clean-up kit (New England Biosciences). Alignment of reads to the human transcriptome (hg38) was performed via `Rsubread` (25).

Cistromes were analyzed with `csaw` (31) with different size windows for transcription factors or histone modifications, along with TF motif analyses `(MotifDb)`. In order to find potential transcription factor binding enrichment within cistromes, `GIGGLE` was utilized to query the complete human transcription factor ChIP-seq dataset collection (10,361 and 10,031 datasets across 1,111 transcription factors and 75 histone marks, respectively) in Cistrome DB(32). Prostate specific filtering limited analysis to 681 datasets across 74 TFs and 238 datasets across 19 HMs. For each query dataset, we determined the overlap of each transcription factor cistrome. Putative co-enriched factors were identified by assessment of the number of time a given factor was observed in the top 200 most enriched datasets relative to the total number of datasets for that factor in the complete Cistrome DB (> 1.2 FC enrichment over background). For prostate specific analysis, overlaps across datasets were averaged for each factor.

### Next generation sequencing

Sequencing was performed at the Nationwide Children’s Hospital Institute for Genomic Medicine, Columbus, OH.

### Data analyses and integration

All analyses were undertaken using the R platform for statistical computing (R version 4.1.3) and the indicated library packages implemented in Bioconductor.

### Data availability

RNA-, ChIP-and ATAC-Seq data are available (***)

## RESULTS

### OSU-ERb-12/Enza inhibits proliferation associated with differentiation

ERβ mRNA and protein expression were measured across a panel of PCa cell lines; non-malignant HPr1AR, androgen responsive PCa, LNCaP, and ADT-R-PCa, LNCaP C4-2, 22Rv1, PC-3 and DU 145 all of which are derived from European American men. We also utilized a series of prostate cells derived from African American (AA) men, namely RC43N and RC77N (non-malignant prostate) and RC43T and RC77T (androgen responsive AA PCa). *ESR2* and ERβ were expressed in all cells, with small but significant elevated mRNA levels in 22Rv1, PC-3 and DU 145 cells, and reduced levels in LNCaP and RC43T cells compared to HPr1AR cells (**Supplementary Figure 1A, B**).

CRISPR approaches were used to generate ERα knockout (ERKO) and ERβ knockout (BERKO) cells confirmed by sequencing (**Supplementary Figure 1C**). BERKO cells had a faster growth rate than parental and ERKO cells (**Supplementary Figure 1D**) and were more invasive in a scratch assay (**Supplementary Figure 1E**) supporting a role for ERβ to inhibit proliferation and limit invasion. Somewhat reflecting the *ESR2*/ERβ expression patterns, OSU-ERb-12 alone was most potent at lower doses such as 0.1 and 1 μM in 22Rv1 and DU145 cells (**Supplementary Figure 1F**), although all cells were significantly inhibited by OSU-ERb-12 at higher doses (10 μM)

Next, we screened dose combinations of OSU-ERb-12 with each of 30 other drugs (**Supplementary Table 1**). Contour plots (**Supplementary Figure 2A**) illustrate in lighter color cooperative inhibition of proliferation between OSU-ERb-12 and the other drugs. For example, there were additive effects in LNCaP, 22Rv1 and PC-3 treated with OSU-ERb-12 plus either Enza or the HDAC inhibitor FK228. We calculated the delta between the observed and predicted values from these data and identified OSU-ERb-12 and drug combinations which resulted in significant additive effects. Positive delta values indicate additive effects between drug combinations, whereas negative delta values indicate antagonism (19) (**Figure 1A, Supplementary Figure 2B**). In LNCaP cells there were additive effects between OSU-ERb-12 and ETYA, an PPARγ agonist; BIQ, an N-6-methyladenosine modulator; and Fisetin, a DNA methyltransferase inhibitor (DMNTi). In PC-3 cells only OSU-ERb-12 plus the HDACi FK228 was additive. DU145 cells responded additively to the combinations of OSU-ERb-12 plus DNMTis including Fisetin or Entinostat, an HDACi. 22Rv1 cells (**Figure 1A**) showed a different set of additive interactions, including with the DNMTi epigallocatechin gallate (EGCG); Enza; and MPP dihydrochloride, an ERα antagonist. Plotting the -log10(p.adj) of the delta values revealed the spectrum additive interactions in each cell, and clustering common events between each cell line and drug combination responses, which supported the common additive combinations of OSU-ERb-12 with either Enza (**Figure 1B**) or FK228 (histone deacetylase inhibitor) (**Supplementary Figure 2B**).

**Figure 1.**
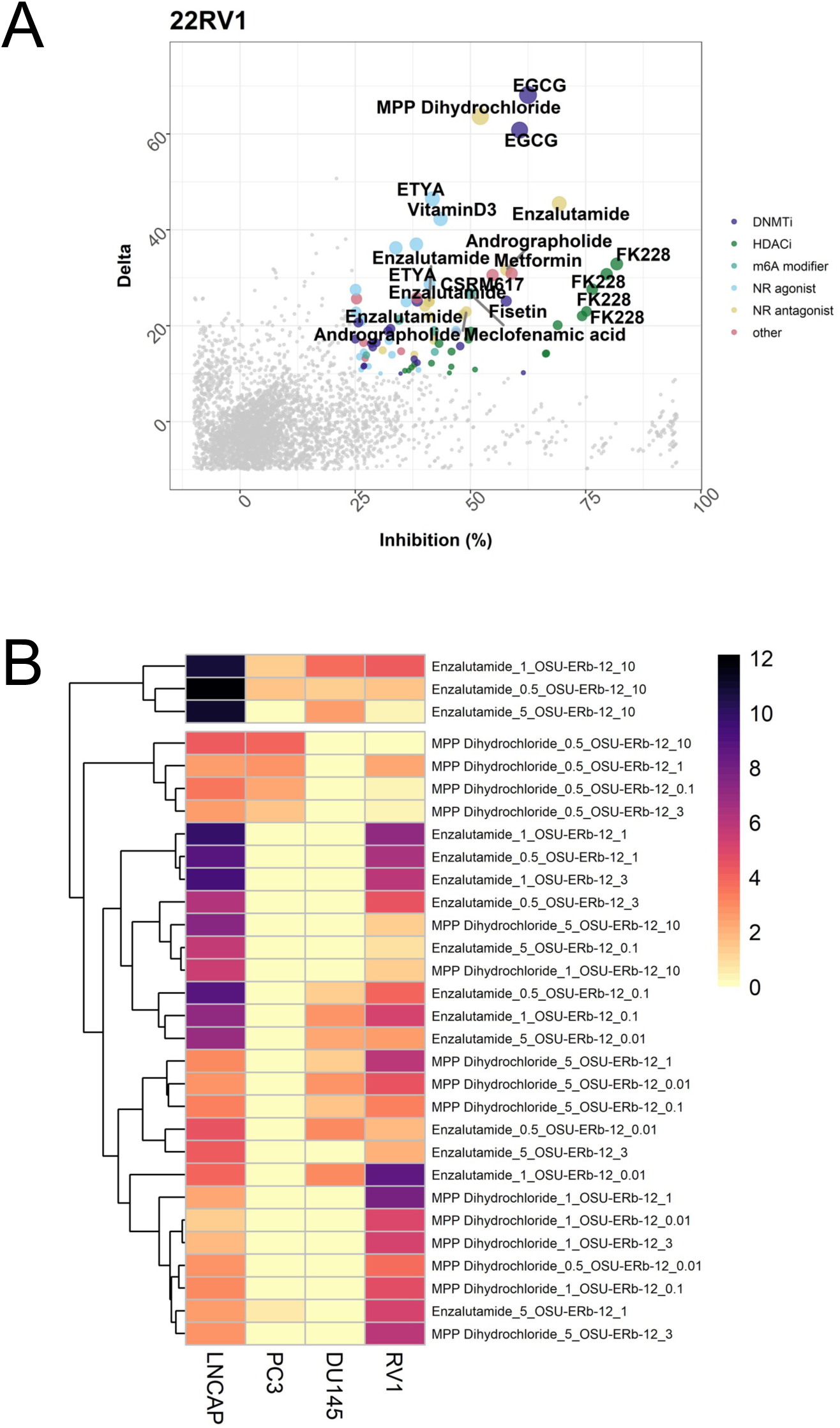
Cooperative actions of OSU-ERb-12 with a range of drugs in LNCaP, PC-3, DU 145 and 22Rv1 cells. **A.** Cells in quadruplicates were treated with a range of OSU-ERb-12 concentrations and a panel of other drugs (Supplementary Table 1) also at a range of doses. Inhibition of proliferation was measured by measuring cellular ATP. The observed combination of proliferation (Inhibition %) was plotted against the difference (delta) to the predicted inhibition; plot for 22Rv1 cells is shown **B.** The difference between the observed and predicted inhibition was measured with a t-test and the p values corrected for multiple testing, log10 transformed and visualized by heatmap. Representative plots are shown for nuclear receptor antagonists.

Clonogenic assays also supported a combinatorial effect of OSU-ERb-12 with Enza (**Figure 2A**), associated with a modest reduction in the levels of programmed cell death (**Supplementary Figure 3**) and sustained expression levels of luminal differentiation markers, namely the AR itself, cytokeratin 5 (CK5), and cytokeratin 8 (CK8) compared to the response to Enza plus 17β-estradiol (E2). Only the combination of OSU-ERb-12 with Enza elevated AR and sustained both CK5 and CK8 levels (**Figure 2B**, **C).**

**Figure 2.**
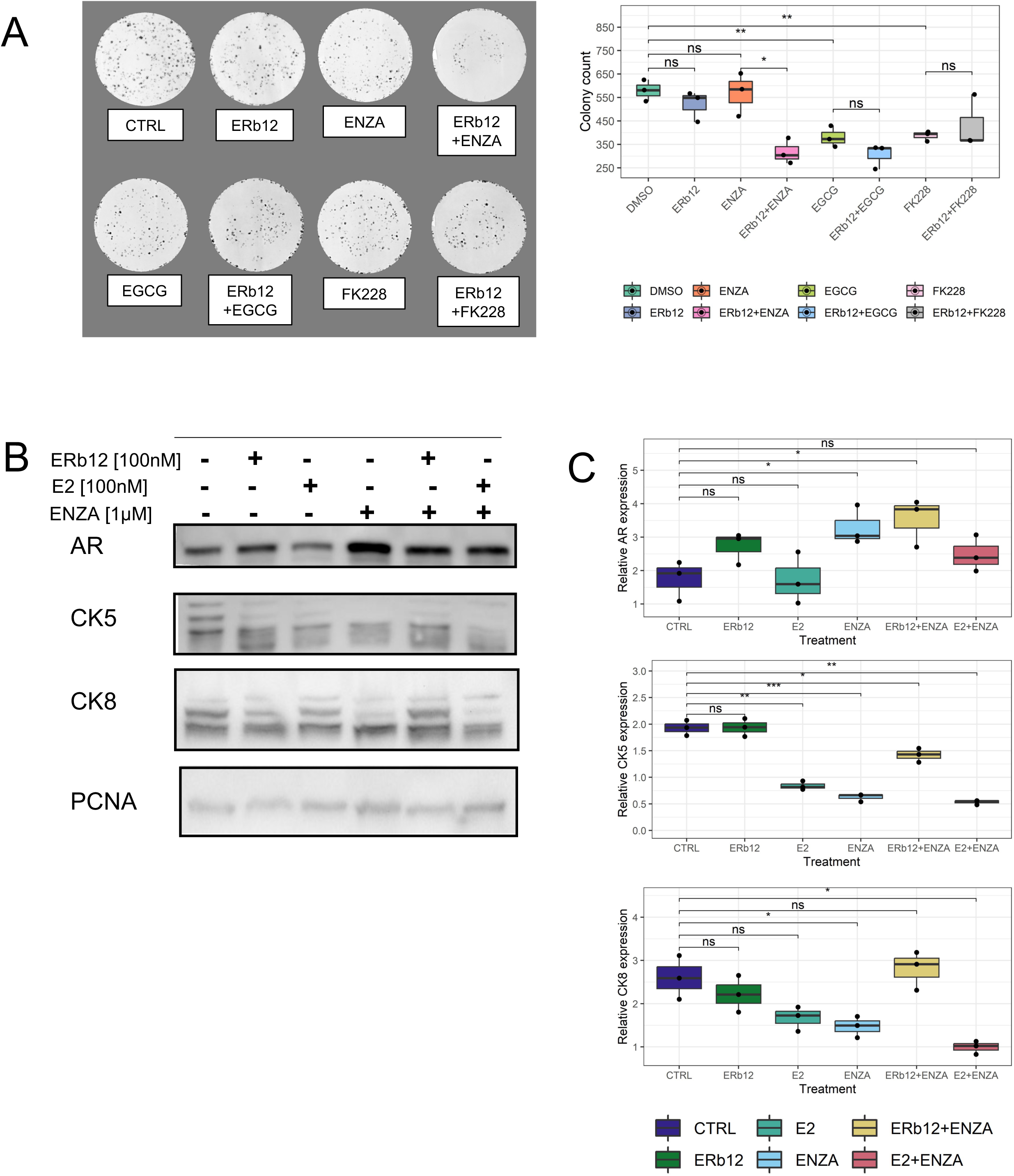
Inhibition of colony formation by OSU-ERb-12 combinations and regulation of differentiation markers in 22Rv1 cells. **A**. Representative images of 22Rv1 colonies treated with OSU-ERb-12 (100 nM) with either Enza (1 μM) or FK228 (1nM) or the combinations with quantification of colony number. **B**. Western immunoblot measurements of AR, CK5 and CK8 after treatment with OSU-ERb-12 plus Enza (time 7 days) with quantification (**C**).

### OSU-ERb-12/Enza represses Androgen and MYC transcriptomes

Q-RT-PCR analyses revealed that OSU-ERb-12 plus ENZA repressed several well-established AR-target genes including TMPRSS2 at 12 h (**Supplementary Figure 4A**). Subsequently, RNA-Seq was undertaken at 12 and 48 h in LNCaP, LNCaP C4-2 and 22Rv1 cells treated with Enza (1 μM). plus either E2 (100 nM) or OSU-ERb-12 (100 nM). Similarity and principal component analyses revealed that experimental conditions explained most of the variation in expression (**Supplementary Figure 4B**).

Differentially expressed genes (DEGs) in response to the individual and combined treatments (**Figure 3A**; DEGs; FDR < 0.1, absolute FC > 1.2) revealed the difference combination treatments of Enza plus E2 plus compared to Enza plus OSU-ERb-12. Specifically, OSU-ERb-12 combinations led to a pronounced gene repression. For example, in LNCaP and LNCaP C4-2 *NKX3.1* was upregulated by E2 plus Enza, whereas this gene was repressed by OSU-ERb-12 plus Enza. Perhaps reflecting how these events are tightly regulated, by 48 h the DEGs were more equally distributed, probably reflecting a combination of primary and secondary effects (**Supplementary Figure 4C**). These findings were complemented by epigenetic landscape *in silico* analysis to measure enrichment of transcription factors (33), which demonstrated that the OSU-ERb-12 plus Enza transcriptome was enriched for genes targeted by transcription factors that promote alternative lineages such as SOX4 and FOXA2 (34,35) as well as MYC and AR (**Supplementary Table 2**).

**Figure 3.**
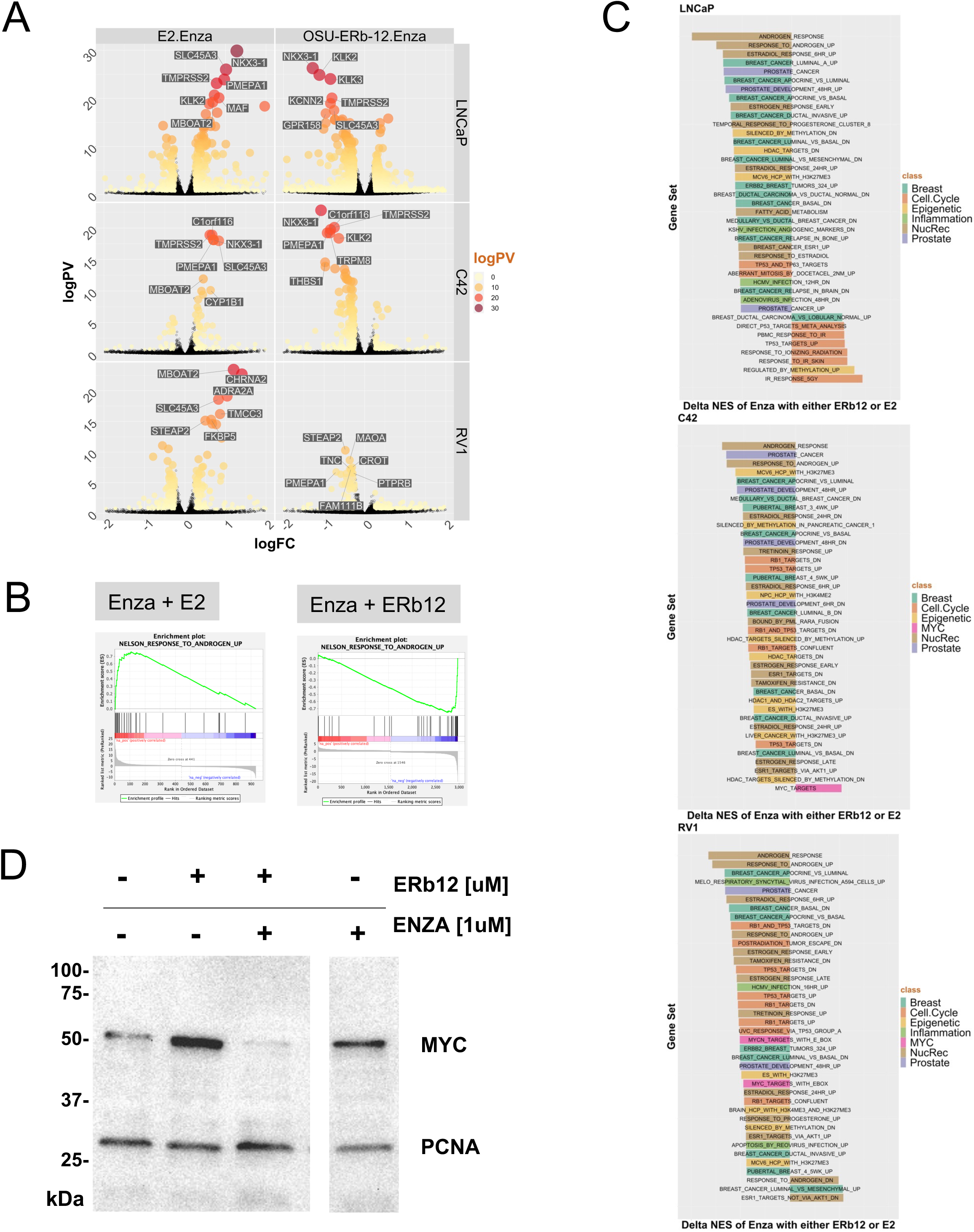
OSU-ERb-12 plus Enza induces a unique transcriptome associated with MYC repression in LNCaP, LNCaP C4-2 and 22 Rv1 cells. **A**. LNCaP, LNCaP C4-2 and 22Rv1 cells in triplicate were treated with either vehicle, OSU-ERb-12 alone (100 nM) or E2 alone (***), Enza (1 μM) or the combinations for 12h and RNA-Seq was undertaken. FASTQ files were QC processed, aligned to hg38 (`Rsubread`), and processed with `edgeR` workflow to identify significant differentially enriched genes (DEGs) (logPV > 1 & absFC > .26) of the drug combination compared to the combination of the average of the individual treatments, and are illustrated by Volcano plots with top DEGs labelled. **B**. GSEA analyses (Hallmarks, and Chemical and Genetic Perturbations) was undertaken and enrichment terms were word frequency analyzed which identified that terms including Breast, Prostate, Nuclear Receptors and MYC were amongst the most frequent, and these terms were filtered for conditions where the NES was in the opposite direction, and the delta calculated, with top 40 NES terms illustrated. **C**. NES plots for the indicated treatments with either OSU-ERb-12 or E2 co-treatments. **D**. Western immunoblot measurements of MYC after treatment with OSU-ERb-12 plus Enza (time 24 h).

LNCaP, LNCAP C4-2 and 22Rv1 cells express four alternatively spliced *ESR2* transcripts, but the most common transcript in all three cells is ENST00000553796.5, which is 1634 aa long and is not altered by OSU-ERb-12 treatments. By contrast, the co-treatment of OSU-ERb-12 plus Enza frequently associated with the greatest number of alternative transcripts. Transcripts were classified(36) as to whether the encoded protein was either a coactivator (CoA), corepressor (CoR), mixed function coregulators (Mixed) or Transcription factors (TF) (**Table 1**). OSU-ERb-12 plus Enza treatment frequently led to more alternative transcripts than E2 plus Enza. For example, OSU-ERb-12 plus Enza in LNCaP led to four CoA DETs including BRD8, whereas E2 plus Enza only led to one CoA DET. This was also clearly seen for the diverse transcripts, classified as “other”, and in LNCaP C4-2.

**Table 1.**
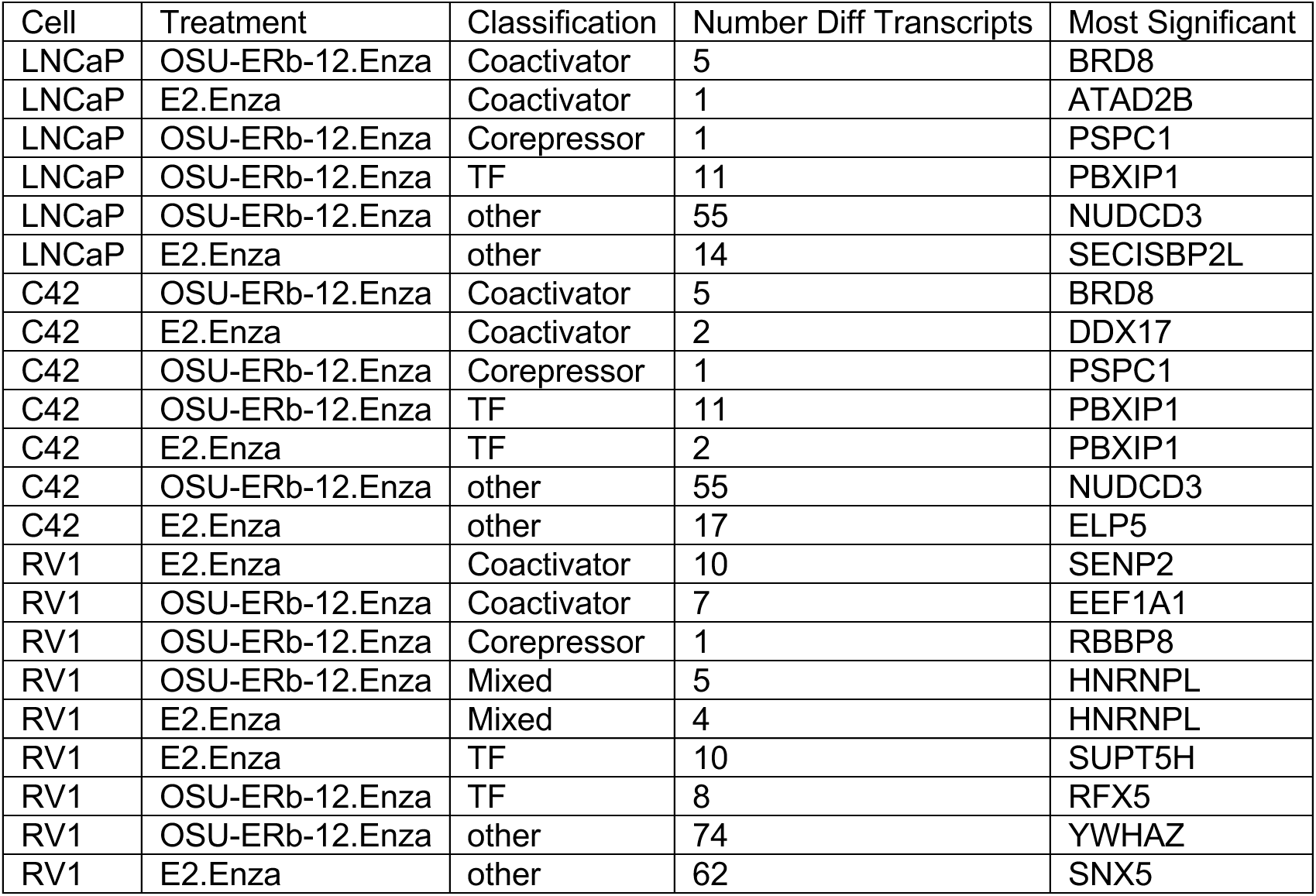
Differentially Expressed Transcripts summarized by treatment and the most significantly altered transcript. LNCaP, C42 and 22Rv1 cells in triplicate were treated with vehicle, Enza (1 μM) plus either E2 (100 nM) or OSU-ERb-12 (100 nM), or the individual treatments and RNA-Seq undertaken. The FASTQ files were aligned with `salmon` and differentially enriched transcripts (DETs) for the combined treatments compared to individual treatments (p.adj < .1) were identified using `DRIMSeq`. Significant DETs were then classified either as a Coactivator (CoA), Corepressor (CoR), Mixed function coregulator (Mixed), transcription factor (TF) or other, and the most significant member of each class in each condition is indicated.

Pre-ranked gene set enrichment analysis (GSEA) was undertaken using the Hallmarks, and Chemical and Genetic Perturbation libraries, and underscored the contrast between E2- dependent activation and OSU-ERb-12 dependent repression from the treatments. We filtered all significant enrichment NES terms (LNCaP ∼800; C42 ∼120; 22Rv1 ∼350) to those that displayed an opposite direction NES score between Enza plus E2 compared to Enza plus OSU-ERb-11, as illustrated for the AR response (**Figure 3B**). We calculated the delta NES score for those terms that switch in NES direction and classified by frequently enriched word terms (breast, prostate, nuclear receptor, and MYC (LNCaP=190; C42=74; 22Rv1=79)) and the top 40 terms are shown in **Figure 3C**. Although nuclear receptor and prostate terms were common across all cells, MYC terms were only enriched in C42 or 22Rv1 cells and was reflected by repression of MYC protein by Enza plus OSU-ERb-12 (**Figure 3D**).

### OSU-ERb-12/ENZA induced nucleosome free regions enriched for basic helix loop helix motifs

To understand further the genomic impact of OSU-ERb-12 plus Enza we undertook ATAC-Seq in 22Rv1 cells at 6h(30). OSU-ERb-12 plus Enza co-treatment led to approximately 3000 unique nucleosome free (NF) regions (**Figure 4A**), which were larger and distinct from the impact of E2 plus Enza (**Supplementary Figure 5A**). Classification of these NF regions in ChromHMM-defined epigenetic states(37) (**Supplementary Table 3**) revealed that OSU-ERb-12 plus ENZA induced NF regions, but not mono-nucleosome regions, were significantly enriched in Active and Poised enhancers, and Polycomb regions (**Supplementary Figure 5B**). These data suggest that the co-treatment induced a uniquely accessible level of chromatin at key regulatory regions.

**Figure 4.**
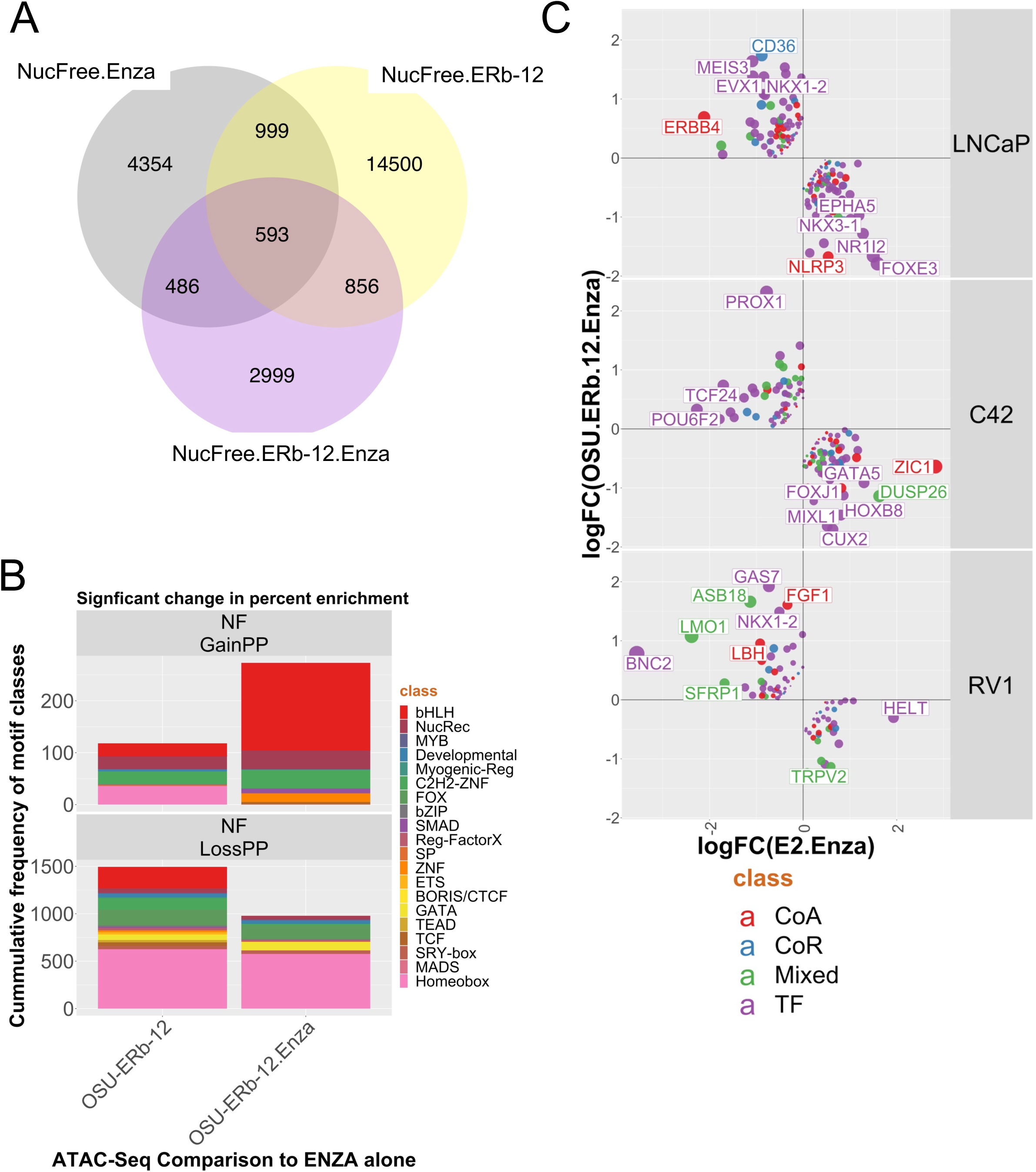
OSU-ERb-12 plus Enza induces distinct patterns of chromatin accessibility in 22Rv1 cells. **A.** 22Rv1 cells were treated with vehicle control, OSU-ERb-12 alone (100 nM), Enza (1 μM) or the combinations for 6 h in triplicate and ATAC-Seq was undertaken. FASTQ files were QC processed, aligned to hg38 (`Rsubread`), sorted and duplicates removed before further processing with `ATACseqQC` to generate nucleosome free (NF) and mono-nucleosome regions. Differential enrichment of regions was measured with `csaw` and the significantly different regions (p.adj < .1) were then intersected to generate the Venn diagrams of overlapping regions by a minimum of 1bp (`ChIPpeakAnno`). **B**. Motif enrichment in NF regions was undertaken with `Homer` and ranked by the percent changes in motifs and those most changing were visualized. **C**. NF regions were annotated to genes (`ChIPpeakAnno`) and those genes overlapped with the transcriptomic data for either OSU-ERb-12 plus Enza, or E2 plus Enza, from which genes were filtered that were regulated in the opposite direction and further filtered to those that were either coactivator (CoA), corepressor (CoR), mixed function coregulators (Mixed) or Transcription factors (TF) and visualized.

Motif enrichment in these unique NF regions was calculated again as the delta for the change in the percentage coverage of motifs between treatment groups compared to Enza alone. To visualize these changes the motifs were then classified by group, for example as nuclear receptors or basic helix loop helix (bHLH) transcription factors, and finally the frequency by group calculated. OSU-ERb-12 plus Enza increased the frequency of coverage of bHLH motifs in NF chromatin (**Figure 4B upper**, red), and there was a reduced representation of homeobox motifs with either OSU-ERb-12 alone or with Enza (**Figure 4B lower**, pink). Within NF chromatin there was also evidence of specific motif enrichment in OSU-ERb-12 plus Enza. For example, with the bHLH factor ASCL1, the nuclear receptor ERE, and the FOXA1:AR motif (**Supplementary Figure 6**).

Integration of the ATAC-Seq and RNA-Seq data also illustrated the antagonism between OSU-ERb-12 and E2 treatments. To illustrate this, we filtered the RNA-Seq data to visualize genes that were **i.** associated with NF chromatin, **ii.** regulated in an antagonistic manner between OSU-ERb-12 and E2 treatments, and **iii.** enriched for coregulators(36). By plotting the OSU-ERb- 12 and E2 combination treatments against each other and filtering these genes for coregulators we identified genes that were oppositely regulated such as *GAS7*(38) a pro-differentiation co-factor is upregulated by OSU-ERb-12 plus Enza but not E2 plus ENZA, as is *LMO1*(39) (**Figure 4C**).

### The combination of OSU-ERb-12 plus Enza leads to unique repression of shared AR-MYC genomic sites

To establish further the genomic impact of OSU-ERb-12 plus Enza we undertook CUT&RUN for AR, ARv7, MYC and H3K27ac. In the first instance, we measured OSU-ERb-12 plus Enza-induced H3K27ac, AR, MYC and ARv7 cistromes, compared to IgG (**Supplementary Figure 7A**). For example, Enza exerted a significant impact on H3K27ac, with many sites shared between control and the other treatments, and indeed ∼2900 H3K27ac sites were constant across all comparisons. Reflecting the ATAC-Seq data, the combination of OSU-ERb-12 plus Enza induced 1880 unique sites of H3K27ac. In control cells there was no significant binding of AR, but ∼1500 ARv7 sites, and the combination of OSU-ERb-12 plus Enza led to ∼8250 AR sites, and ∼1250 ARv7 sites. The co-treatment also led to ∼6200 unique MYC binding sites (**Supplementary Figure 7B**).

To define the individual and combined treatments we focused on H3K27ac cistromes (**Supplementary Figure 7A**) and annotated the 15 individual sub-sets in this Venn diagram to genes. Within these genes we again examined enrichment of coregulators by a hypergeometric test (**Figure 5A**), which revealed that the 1880 unique H3K27ac sites (ERb12Enza) were significantly enriched for all classes of coregulators, TFs and also NRs as a class, with *ESRRG* as the closest NR. Indeed, the levels of enrichment were comparable to those measured for the ∼6800 sites shared between OSU-ERb-12 PLUS Enza and Enza alone (Enza.ERb12Enza), suggesting that the unique aspects of OSU-ERb-12/Enza treatment were targeted to gene rich regions associated with coregulators.

**Figure 5.**
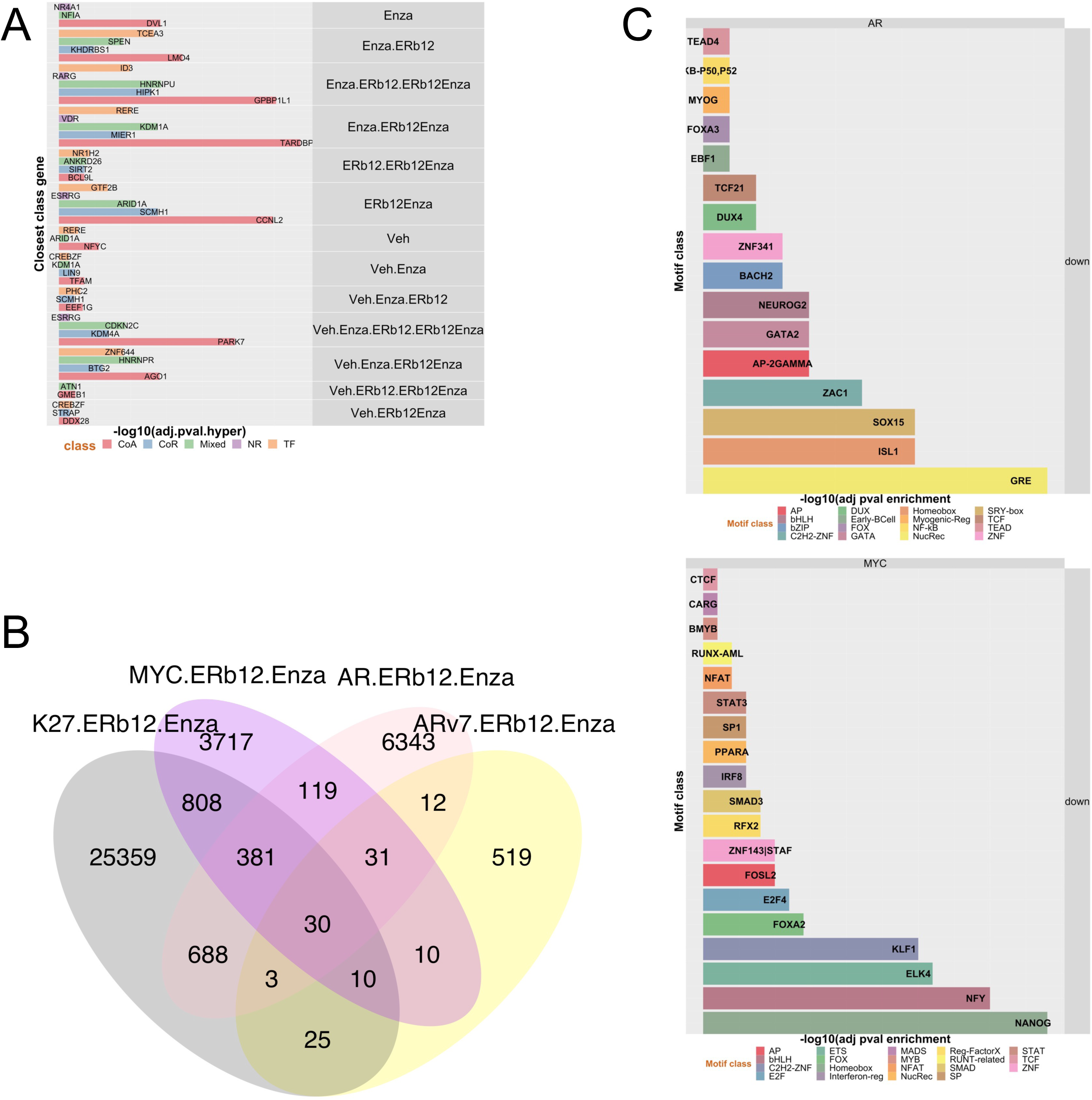
AR, ARv7, MYC and H3K27ac cistromes respond to OSU-ERb-12 plus Enza. 22Rv1 cells were treated with vehicle control, OSU-ERb-12 alone (100 nM), Enza (1 μM) or the combinations for 6 h in triplicate and CUT&RUN undertaken with antibodies to AR, ARv7, MYC or H3K27ac. FASTQ files were QC processed, aligned to hg38 (`Rsubread`), sorted and duplicates and differential enrichment of regions of the drug combination compared to the average of the individual treatments with `csaw`. **A.** The significant differentialy enriched regions, both up and down) in the combined treatments for the AR, ARv7, MYC and H3K27ac cistromes were intersected to generate the Venn diagram of overlapping regions by a minimum of 1bp (`ChIPpeakAnno`). **B.** The H3K27ac cistromes of the individual and combined treatments, compared to IgG, were intersected (minimum of 1bp) and the genomic regions in each intersection (**Supplementary Figure 7A**) were annotated to genes within 100 kb, and filtered as to whether the genes were either coactivator (CoA), corepressor (CoR), mixed function coregulators (Mixed) or Transcription factors (TF). Enrichment was then determined by hypergeometric test and the group gene member closest to a given H3K27ac site is indicated. **C.** Motif analyses of MYC and AR sites that were significantly reduced or lost by the cotreatment and the most significant of which is illustrated by transcription class terms and the most significant class member indicated.

We also measured the overlap of the significantly distinct OSU-ERb-12 plus Enza cistromes (calculated compared to the individual treatments) (**Figure 5B**). For example, this revealed that the co-treatment led to a large increase in unique AR binding sites (∼6350 sites) and that the most significant overlaps were between AR and MYC sites (p=3.1e^-271^), followed by AR and H3K27ac (3.2e^-160^), and then ARv7 and MYC (2.4e^-57^). Next, we annotated these cistrome data to genes by dividing sites into those that were with gained or lost. Considering the pronounced transcriptomic impact of OSU-ERb-12 plus Enza on gene repression, many of the sites impacted by the cotreatment of OSU-ERb-12 plus Enza led to reduced enrichment for AR, MYC, and H3K27ac (**Supplementary Figure 8**). For example, the cotreatment reduced MYC and AR binding at ∼ 3990 and ∼6500 MYC and AR sites respectively, compared to the individual treatments alone, and there was a small but significant overlap MYC and AR (∼220 sites, p=1.6e^-136^). Further annotation of all AR, MYC, and H3K27ac sites to ChromHMM regions revealed that the sites of reduced MYC, AR and H3K27ac binding were enriched in poised enhancers and active enhancers (**Supplementary Table 2**). This was supported further with motif enrichment analyses which revealed that sites of AR binding loss were highly enriched for nuclear receptors (e.g. glucocorticoid receptor), Homeobox (e.g. ISL1) and SRY-box (e.g. SOX15), and the sites of MYC binding loss were enriched for Homeobox also (e.g. NANOG), MADS (e.g. NFY) and ETS (e.g. ELK4) (**Figure 5C**). AR, MYC and H3K27ac binding sites that displayed gain or loss were annotated to genes and then tested for the enrichment of coregulators (**Table 2**). Annotation to of the individual and shared regions of altered enrichment to genes revealed that gained H3K27ac and MYC sites were significantly enriched for coactivators including ZBTB48(40). Similar annotation of regions of AR, MYC and H3K27ac regions revealed enrichment around corepressors including RNF2(41) and KDM4A which is a repressor of MYC(42). Interestingly, there was significant loss of AR/H3K27 and AR/MYC enrichment at nuclear receptors as a class, including ESRRG, which is a known AR target(43) (**Supplementary Table 4**).

**Table 2:** Significant enrichment of AR, MYC or H3K27ac cistromes in ChromHMM defined epigenetic states. 22Rv1 cells in triplicate were treated with vehicle, Enza (1 μM) plus OSU-ERb- 12 (100 nM), or the individual treatments and CUT&RUN undertaken for AR, MYC or H3K27ac. Differential enriched binding sites for the combined treatments compared to individual treatments (p.adj < .1) were identified by `csaw` and were overlapped using `bedtools` with ChromHMM regions identified in LNCaP, and enrichment tested with a hypergeometric test (lower.tail = FALSE).

| ChromHMM | ChIP | Enrichment | logPval | Threshold |
| --- | --- | --- | --- | --- |
| Poised_Enhancer | MYC | loss | 188.66 | Significant |
| Poised_Enhancer | K27 | loss | 174.98 | Significant |
| Poised_Enhancer | AR | loss | 169.30 | Significant |
| Polycomb | K27 | loss | 90.29 | Significant |
| Polycomb | K27 | gain | 72.13 | Significant |
| Poised_Enhancer | K27 | gain | 41.58 | Significant |
| Active_Enhancer | K27 | loss | 27.36 | Significant |
| Active_Enhancer | MYC | loss | 25.96 | Significant |
| Active_Enhancer | AR | loss | 21.97 | Significant |
| Promoter | K27 | loss | 16.40 | Significant |
| Bivalent_Promoter | AR | loss | 10.37 | Significant |
| Bivalent_Promoter | K27 | loss | 6.64 | Significant |
| Poised_Enhancer | MYC | gain | 4.63 | Significant |
| Active_Enhancer | K27 | gain | 3.16 | Significant |
| Bivalent_Promoter | MYC | loss | 2.87 | Significant |
| Bivalent_Promoter | K27 | gain | 2.02 | Significant |
| Polycomb | MYC | loss | 1.20 | Significant |

To understand these cistromes more fully we undertook comprehensive cistrome enrichment analyses with GIGGLE(32) to identify how the AR and MYC cistromes overlapped with TF and histone modifications contained in the CistromeDB collection (> 10,000 total ChIP-seq datasets, across > 1100 factors). Focusing on the regions where OSU-ERb-12 plus Enza led to loss of AR or MYC binding compared to agent alone revealed how these cistromes significantly overlapped with a large number of other cistromes all derived in PCa cell lines also. Reassuringly the AR cistrome overlapped with AR cistromes, as well as ERα, GR and VDR cistromes and the ETS member ERF. Generally, the MYC cistrome was more enriched for the same factors, and included other MYC cistromes, and lineage regulators such as ONECUT2, NANOG, HOXB13, and BRD4 (**Figure 6A**).

**Figure 6.**
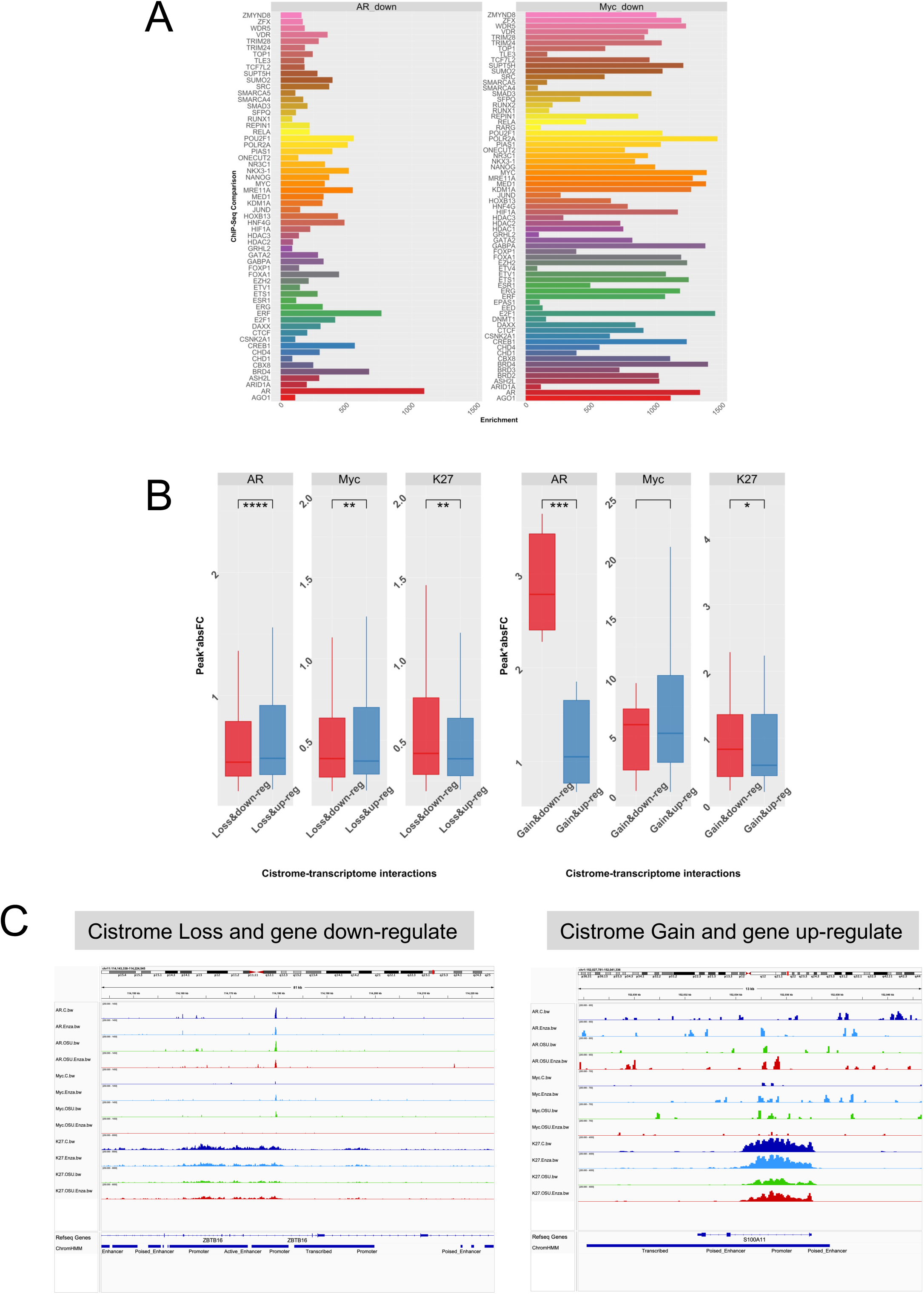
Cistrome-transciptome relationships impacted by OSU-ERb-12 plus Enza. **A**. Unique AR and MYC cistromes in response to OSU-ERb-12/Enza were overlapped using GIGGLE with the CistromeDB collection (> 10,000 total ChIP-seq datasets, across > 1100 factors). The logPV of the FDR-corrected pval is illustrated for prostate specific TFs and coregulators. **B.** Each peak was annotated to genes within 100 kb and filtered as to whether the gene was a DEG from the respective RNA-Seq and the score.absFC calculated for peak:gene relationships that were also DEGs, and a t-test used to assess whether these values were significantly different between the groups where the CUT&RUN peak enrichment was reduced (D) or increased (U) and the annotated gene expression what down-regulated (D) or up-regulated (U). **C.** IGV image of the AR, MYC and H3K27ac at the ZBTB16 gene. Triplicate BAM files were merged and converted to bigwig files and visualized at the ZBTB16 gene as an example of a gene that displays significantly reduced enrichment by OSU-ERb-12/Enza for H3K27ac and MYC, and is downregulated at the mRNA level.

Finally, these OSU-ERb-12/Enza differential H3K27ac, AR and MYC binding sites were integrated with the RNA-Seq data for the same conditions. The cistrome-transcriptome relationships were classified by the direction of the ChIP enrichment (Loss or Gain) and RNA expression (down-or upregulation) and the peak score/gene, calculated as the peak score multiplied by the absolute Fold Change for the same gene (score.absFC)(44). This revealed that cotreatment of OSU-ERb-12 plus Enza significantly impacted AR, MYC and H3K27 relationships to gene expression (**Figure 6B**). The score.absFC values were modestly but significantly greater for genes that demonstrated a loss of AR binding in response to OSU-ERb-12 plus Enza and were upregulated rather than downregulated, suggesting the co-treatment overcomes intrinsic AR repression of these genes. By contrast, genes where OSU-ERb-12 plus Enza reduced MYC and H3K27ac enrichment were more strongly downregulated than upregulated. A smaller number of genes were identified where OSU-ERb-12 plus Enza increased AR binding and they were significantly more downregulated than up regulated, and this was also seen with H3K27ac. ZBTB16 and S100A11 illustrated these cistrome-transcriptome relationships.

Finally, in the SU2C cohort(45), we examined expression of 192 genes from the above cistrome-transcriptome analyses, specifically, those genes with loss of AR enrichment and upregulation, and loss of MYC and H3K27ac enrichment and gene downregulation. These genes significantly separated tumors by AR score (Pearson’s Chi-squared test with Yates’ continuity correction; X^2^=81.2; pval = 2.2e^-16^) and neuroendocrine score (X^2^=15.3; p-value = 0.00048) (**Figure 7**) and supported the concept that OSU-ERb-12 plus Enza uniquely impacts AR, MYC and H3K27ac cistromes in a manner that regulates genes that in turn are significantly associated with sustained AR status and impede acquisition of alternative lineages such as NE-PCa.

**Figure 7.**
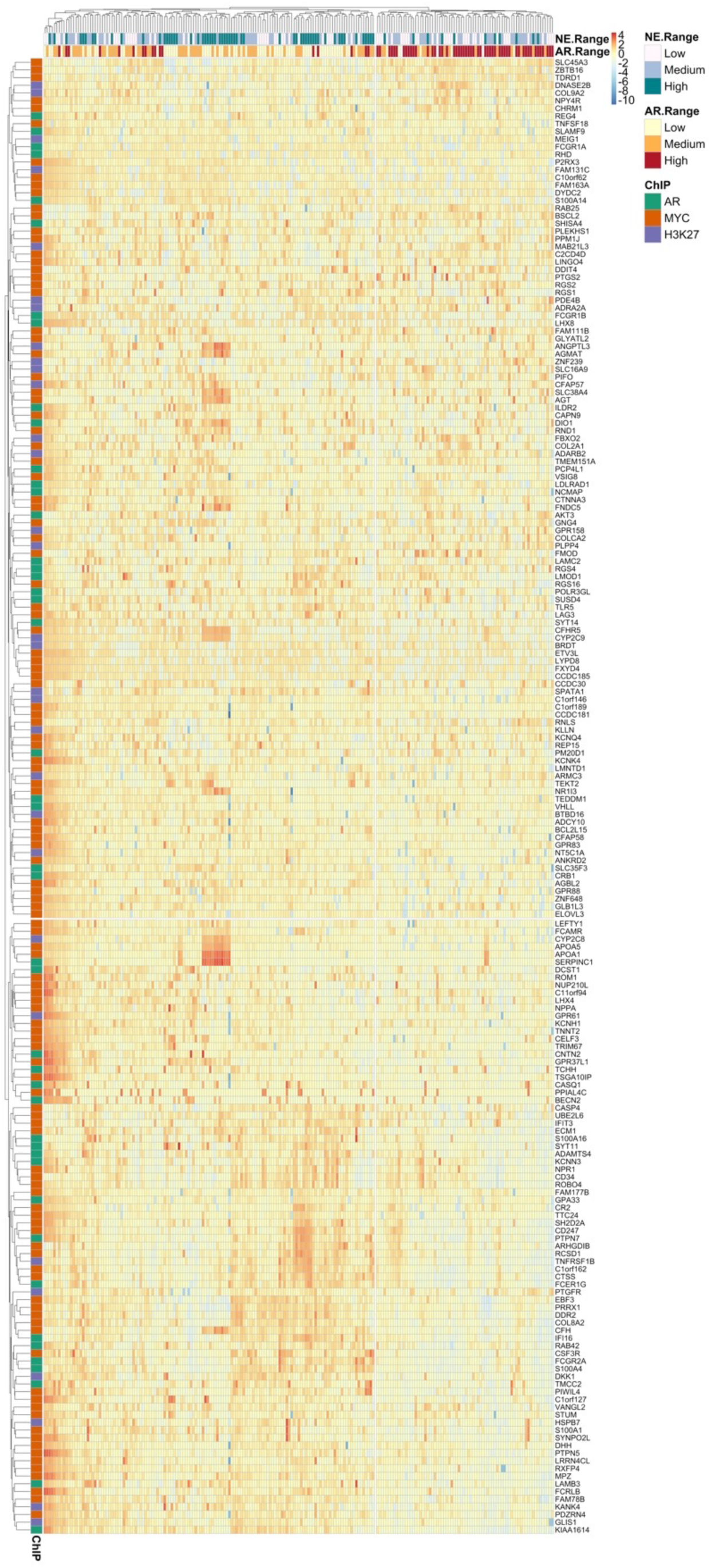
Unique OSU-ERb-12 plus Enza cistrome-transciptome dependent target genes significantly associate with high androgen and low neuroendocine score tumors in the SU2C cohort. The Zscore data frame from the SU2C cohort was filtered for 192 unique genes that were defined by; a loss of AR binding and upregulation by OSU-ERb-12 plus Enza; a loss of MYC binding and downregulation by OSU-ERb-12 plus Enza; a loss of H3K27ac binding and downregulation by OSU-ERb-12 plus Enza. These relationships were visualized with a heatmap (`pheatmap`) and tumors clustered by expression, and the significance of the relationship between tumor cluster id and either neuroendocrine score (NE range) or androgen score (AR range), as define previously(45), was measured with a chi-sq test.

## DISCUSSION

In the current study we set out to examine how a selective ERβ with strong pharmacodynamic properties(16,46) could be combined with existing therapies to enhance potency further, and what might be the genomic impact. Supportively, ERβ is expressed at comparable levels across prostate cell models, and it regulates proliferation and invasiveness including in ADT-recurrent models such as 22Rv1 cells. Initial experiments focused on how to enhance OSU-ERb-12 potency. Across four PCa cell lines, including three ADT-R-PCa models, the greatest inhibition was observed with combinations of OSU-ERb-12 with either Enza, or the histone deacetylase inhibitor, FK228, or EGCG, a DNMT inhibitor; these latter two compounds suggesting a common role for epigenetic events to suppress ERβ functions. Phenotypic analyses revealed that OSU-ERb-12 plus Enza demonstrated significant cooperation on the inhibition of cell growth and colony formation as well as some evidence that this process was controlling differentiation. Generally, the combination of OSU-ERb-12 plus Enza was more potent than the comparison of E2 plus Enza.

Across LNCaP, LNCaP C4-2 and 22Rv1 cells transcriptomic analyses OSU-ERb-12 plus Enza revealed a global skew to repression of gene expression compared to E2 plus Enza which skewed to activation. Notably, some of the most significant gene expression changes were for AR target genes such as *NKX3*.1 and *TMPRSS2*, and indeed GSEA analyses supported the switch between activation to repression by OSU-ERb-12 plus Enza of AR gene signatures. Indeed, gene set enrichment analyses revealed a pronounced switching of NES scores between OSU-ERb-12 compared to E2 treatments including for terms related to epigenetic events, prostate and cell cycle. Of note in 22Rv1 cells this included repression of MYC terms and repression of protein level. Similarly, epigenomic analyses revealed enrichment of MYC as a driver transcription factor in the genes regulated by OSU-ERb-12 plus Enza. Isoform-aware alignment also revealed that OSU-ERb-12 plus Enza induced high levels of alternative splicing, including for a number of coregulators, and reflects a known function for ERβ to associate with AGO2 and control splice variation(47).

ATAC-seq experiments demonstrated that OSU-ERb-12 plus Enza also uniquely increased nucleosome free chromatin regions that were enriched for enhancers. Motif enrichment revealed that the co-treatment led to a pronounced increase in bHLH motif, which reflects the enrichment of MYC signaling in the transcriptomic data(48), as well as changes to AR half-sites motifs. These data were integrated with the RNA-Seq data, and further demonstrated the antagonistic impact between the two combinations. Again, filtering ATAC-Seq annotated genes for those regulated by either OSU-ERb-12 plus Enza or E2 plus Enza identified ∼400 transcription factors and coregulators where OSU-ERb-12 plus Enza uniquely changed chromatin accessibility and oppositely-regulated gene expression, including *ZBTB16/PLZF*(49).

We also mined the impact of OSU-ERb-12 plus Enza on the AR, ARv7, MYC and H3K27ac cistromes, which further demonstrated a unique impact of the cotreatment compared to either agent alone. CUT&RUN demonstrated a significant loss of enrichment of H3K27ac, AR, and MYC cistromes, reflecting the transcriptomic data, and again that the changes in enrichment were significantly enriched in ChromHMM-identified regulatory regions including active enhancers, there. There was also significant overlap between these cistromes, which was impacted OSU-ERb-12 plus Enza. For example, in the basal state ARv7 cistrome was strikingly larger than AR, but the co-treatment elevated AR, whereas ARv7 was approximately constant. Analyses of the H3K27ac cistrome revealed that the cotreatment significantly impacted regulatory regions associated with coregulators. Focusing on regions where the co-treatment reduced enrichment revealed ∼220 sites of shared AR and MYC binding, and more broadly these regions of reduced binding were themselves enriched for nuclear receptor motifs including the GR in the case of the AR, and the bHLH factor NFY in the case of MYC, and these cistromes significantly overlapped with multiple PCa-specific cistromes. Overlap with the RNA-seq data, again revealed that enrichment in enhancer regions and also there was significant overlap also between where AR and MYC were binding and genes being regulated, and supported a role for OSU-ERb-12 plus Enza to shape the relationship between AR and MYC binding and their relationship to gene expression, for example with *ZBTB16/PLZF*(49). Indeed, cistrome-transcriptome analyses identified ∼200 genes that significantly distinguished AR and NE-PCa in the SU2C cohort of advanced PCa.

These data support a role for OSU-ERb-12 plus Enza to control chromatin accessibility around enhancers and regulatory regions that are co-targeted by the AR and MYC, and selectively-transrepresses AR signaling. This impact on AR and MYC signaling in advanced PCa occurs in a manner that is distinct from E2 plus Enza and illuminates the distinctive biology between ERα and ERβ. These findings are also in keeping with recent studies identifying roles for AR repression of MYC(50), and suggest that there is pivotal role for the ERβ to sustain AR-dependent phenotypes, which can be targeted pharmacologically, and the footprint genes of this regulation are able to distinguish advanced PCa with high AR and low NE scores.

## ABBREVIATIONS

ADT: Androgen Deprivation Therapy
AR: Androgen receptor
ATAC-Seq: Assay for Transposase-Accessible Chromatin using sequencing
CUT&RUN: Cleavage Under Targets and Release Using Nuclease
DHT: Diydroxytestosterone
MYC: MYC Proto-Oncogene, BHLH Transcription Factor
NE-PCa: Neuroendocrine-Prostate cancer
RB: RB Transcriptional Corepressor 1
RNA-Seq: next generation sequencing of RNA
TCGA: The Cancer Genome Atlas
TCGA-SU2C: Stand up to Cancer cohort in the TCGA
TF: Transcription factor

## DECLARATIONS

### Authors’ contributions

*JSG* undertook CUT&RUN, RNA-Seq, ATAC-Seq and cell biology; *MDL* undertook GIGGLE analyses; *CD, CCC, SKC* contributed to experimental design and contributed to data interpretation; *CCC, CD* and *MJC* jointly conceived of the study design; *MJC* oversaw the implementation of the study and *MJC* undertook bioinformatic analyses and generated tables and figures.

## Acknowledgement

Dr. Werner Tjarks is gratefully acknowledged for his innovative development of the chemical synthesis of para-carborane ERβ-selective agonists such as OSU-ERb-12.

