## SupplementaryTablesFigures for "The MYC axis in advanced prostate cancer is impacted through concurrent targeting of ERβ and AR using a novel ERβ-selective ligand alongside Enzalutamide"

Supplementary tables
Supplementary figure legends

| Compound | Class | Function | Low concentration | Medium concentration | High concentration |
|---|---|---|---|---|---|
| OSU-ERb-12 | | ERβ-selective ligand | | | |
| Epigallocatechin gallate (EGCG) | DNMT inhibitor | non-nucleoside inhibitor | 5 | 10 | 20 |
| RG108 | DNMT inhibitor | non-nucleoside inhibitor | 5 | 10 | 20 |
| Decitabine | DNMT inhibitor | nucleoside inhibitor | 0.1 | 0.5 | 1 |
| Fisetin | DNMT inhibitor | nucleoside inhibitor | 1 | 10 | 20 |
| GSK126 | DNMT inhibitor | nucleoside inhibitor | 0.5 | 1 | 10 |
| Hydralazine Hydrochloride | DNMT inhibitor | nucleoside inhibitor | 5 | 10 | 20 |
| Zebularine | DNMT inhibitor | nucleoside inhibitor | 5 | 10 | 20 |
| FK228 | HDAC inhibitor | general HDAC inhibitor | 0.1 | 0.5 | 1 |
| Entinostat | HDAC inhibitor | HDAC 1 and 3 inhibitor | 0.1 | 0.5 | 1 |
| Vorinostat | HDAC inhibitor | HDAC 1 and 2 inhibitor | 0.1 | 0.5 | 1 |
| Enzalutamide | NR antagonist | Androgen receptor antagonist | 0.5 | 1 | 5 |
| MPP Dihydrochloride | NR antagonist | Estrogen receptor alpha antagonist | 0.5 | 1 | 5 |
| All trans retinoic acid (ATRA) | NR agonist | Retinoic acid receptor (RAR) agonist | 0.1 | 0.5 | 1 |
| CD437 | NR agonist | RAR? agonist | 0.1 | 0.5 | 1 |
| ST1926 | NR agonist | RAR agonist | 0.1 | 0.5 | 1 |
| Eicosatetraynoic acid (ETYA) | NR agonist | Peroxisome proliferator-activated receptor gamma (PPARγ) agonist | 1 | 5 | 10 |
| Bezafibrate | NR agonist | PPARα agonist | 5 | 10 | 20 |
| Vitamin D3 | NR agonist | Vitamin D receptor agonist | 0.01 | 0.05 | 0.1 |
| AS1842856 | transcription factor inhibitor | FOXO1 inhibitor | 0.1 | 0.5 | 1 |
| CSRM617 | transcription factor inhibitor | ONECUT2 inhibitor | 0.5 | 1 | 5 |
| GN25 | transcription factor inhibitor | Snail inhibitor | 0.1 | 0.5 | 1 |
| Melatonin | transcription factor inhibitor | Inhibitor of OCT4 | 0.1 | 0.5 | 1 |
| BIQ | m6A modulator | Hypoxia inducible factor alpha inhibitor | 0.1 | 0.5 | 1 |
| Citrate | m6A modulator | AlkB homolog 5 inhibitor | 10 | 50 | 100 |
| Meclofenamic acid | m6A modulator | FTO protein inhibitor | 0.5 | 1 | 5 |
| Sinefungin | m6A modulator | S-adenosyl methionine analog | 0.1 | 0.5 | 1 |
| 2D08 | other | inhibitor of UBC9 (sumoylation) | 1 | 5 | 10 |
| Andrographolide | other | NFκB inhibitor | 1 | 5 | 10 |
| Metformin | other | potential inhibitor of epithelial to mesenchymal transition | 12500 | 25000 | 50000 |
| PFI1 | other | BET bromodomain inhibitor BRD2/4 | 0.1 | 0.5 | 1 |

**Supplementary Table 1.** Drugs used in the single and combined proliferation studies in a panel of prostate cancer cell lines.

| Transcription.Factor | rank | Enrichment (p.val) |
|---|---|---|
| SOX4 | 1 | 1.93e-15 |
| ZBTB48 | 2 | 2.84e-15 |
| FOXA2 | 3 | 5.16e-15 |
| MYC | 4 | 4.9e-12 |
| AFF1 | 5 | 5.83e-12 |
| FOXA1 | 6 | 6.11e-12 |
| MBOAT4 | 7 | 6.7e-12 |
| AR | 8 | 1.5e-11 |
| PR | 9 | 2.08e-11 |
| CEBPB | 10 | 3.35e-11 |
| ERCC2 | 11 | 6.05e-11 |
| AGO1 | 12 | 7.84e-11 |
| HDAC1 | 13 | 1.82e-10 |
| LMNB1 | 14 | 1.89e-10 |
| XRN2 | 15 | 2.76e-10 |

**Supplementary Table 2**: Epigenetic landscape *in silico* analysis of DEGs. RNA-Seq data was analyzed with LISA to identify commonly enriched transcription factors.

| ChromHMM | nuc | RX | logPval | Threshold |
|---|---|---|---|---|
| Poised_Enhancer | NF | E2.Enza | 299.22 | Significant |
| Poised_Enhancer | mono | OSU-ERb-12 | 299.22 | Significant |
| Poised_Enhancer | NF | OSU-ERb-12 | 299.22 | Significant |
| Poised_Enhancer | NF | Enza | 299.22 | Significant |
| Polycomb | mono | OSU-ERb-12 | 251.81 | Significant |
| Polycomb | NF | Enza | 231.15 | Significant |
| Poised_Enhancer | NF | OSU-ERb-12.Enza | 221.97 | Significant |
| Poised_Enhancer | mono | Enza | 98.47 | Significant |
| Polycomb | NF | OSU-ERb-12.Enza | 88.18 | Significant |
| Poised_Enhancer | mono | E2.Enza | 79.36 | Significant |
| Polycomb | mono | Enza | 75.98 | Significant |
| Polycomb | mono | E2.Enza | 50.55 | Significant |
| Transcribed | mono | OSU-ERb-12 | 41.10 | Significant |
| Active_Enhancer | NF | E2.Enza | 40.42 | Significant |
| Active_Enhancer | NF | OSU-ERb-12 | 40.42 | Significant |
| Polycomb | NF | E2.Enza | 38.39 | Significant |
| Polycomb | NF | OSU-ERb-12 | 38.39 | Significant |
| Active_Enhancer | mono | OSU-ERb-12 | 29.84 | Significant |
| Active_Enhancer | NF | OSU-ERb-12.Enza | 27.18 | Significant |
| Transcribed | NF | Enza | 19.61 | Significant |
| Active_Enhancer | mono | E2.Enza | 13.34 | Significant |
| Transcribed | mono | Enza | 12.82 | Significant |
| Transcribed | mono | E2.Enza | 6.39 | Significant |
| Active_Enhancer | mono | Enza | 5.41 | Significant |
| Active_Enhancer | NF | Enza | 4.37 | Significant |

**Supplementary Table 3**: Significant enrichment of drug-induced chromatin accessibility cistromes in ChromHMM defined epigenetic states. 22Rv1, cells in triplicate were treated with vehicle, Enza (1 $\mu$M) plus OSU-ERb-12 (100 nM), or the individual treatments, ATAC-Seq undertaken and nucleosome spacing determined (nucleosome free (NF), or mono-nucleosome (mono) and differential enriched binding sites for the combined treatments compared to individual treatments (p.adj < .1) were identified by `csaw` and were overlapped using `bedtools` with ChromHMM regions identified in LNCaP, and enrichment tested with a hypergeometric test (lower.tail = FALSE).

| ChIP Target | Direction | Nearest Gene | Class | logPV.h |
|---|---|---|---|---|
| MYC.K27 | Gain | ZBTB48 | CoA | 3.48 |
| MYC.K27 | Gain | PHF13, MIB2 | Mixed | 3.21 |
| MYC.K27 | Gain | | Mixed | 3.21 |
| MYC | Gain | CAMTA1, GABPB2 | CoA | 1.52 |
| MYC | Gain | | CoA | 1.52 |
| MYC | Gain | CHD5 | Mixed | 1.41 |
| AR | Gain | TBX19 | CoA | 1.33 |

| ChIP Target | Direction | Nearest Gene | Class | logPV.h |
|---|---|---|---|---|
| AR.K27 | Loss | ESRRG | NR | 3.56 |
| AR.K27 | Loss | RNF2 | CoR | 1.19 |
| AR.MYC | Loss | PMF1, RBBP5 | CoA | 1.02 |
| AR.MYC | Loss | | CoA | 1.02 |
| AR.MYC | Loss | KDM4A, OTUD7B, CIART, SETDB1, MDM4 | CoR | 1.0 |
| AR.MYC | Loss | | CoR | 1.0 |
| AR.MYC | Loss | | CoR | 1.0 |
| AR.MYC | Loss | | CoR | 1.0 |
| AR.MYC | Loss | | CoR | 1.0 |
| AR.MYC | Loss | RXRG | NR | 1.0 |

**Supplementary Table 4**: Enrichment of coregulators and nuclear receptors in AR, MYC and H3K27ac cistromes in response to ERb12 plus Enza. The indicated cistromes following ERb12 plus Enza compared to either treatment alone were identified by `csaw`, and separated into sites that were gained (upper) or lost (lower), the sites were intersected (minimum of 1bp) and the genomic regions in each intersection annotated to genes within 100 kb, and filtered as to whether the genes were either coactivator (CoA), corepressor (CoR), mixed function coregulators (Mixed), Transcription factors (TF) or Nuclear Receptor (NR). Enrichment was then determined by hypergeometric test and the group gene member closest to a region is indicated (more than one indicated is for regions at the TSS).

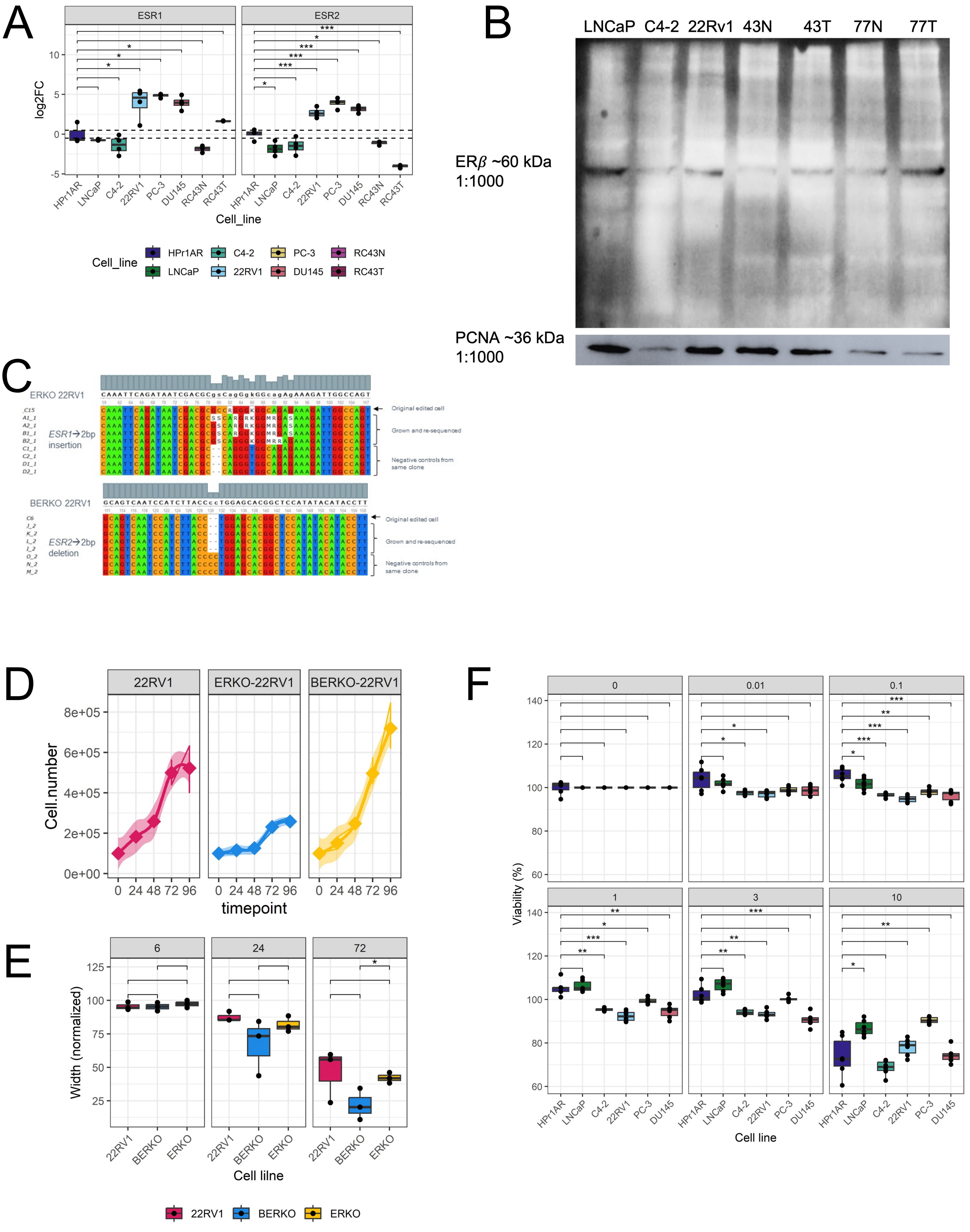

Supplementary Figure 1

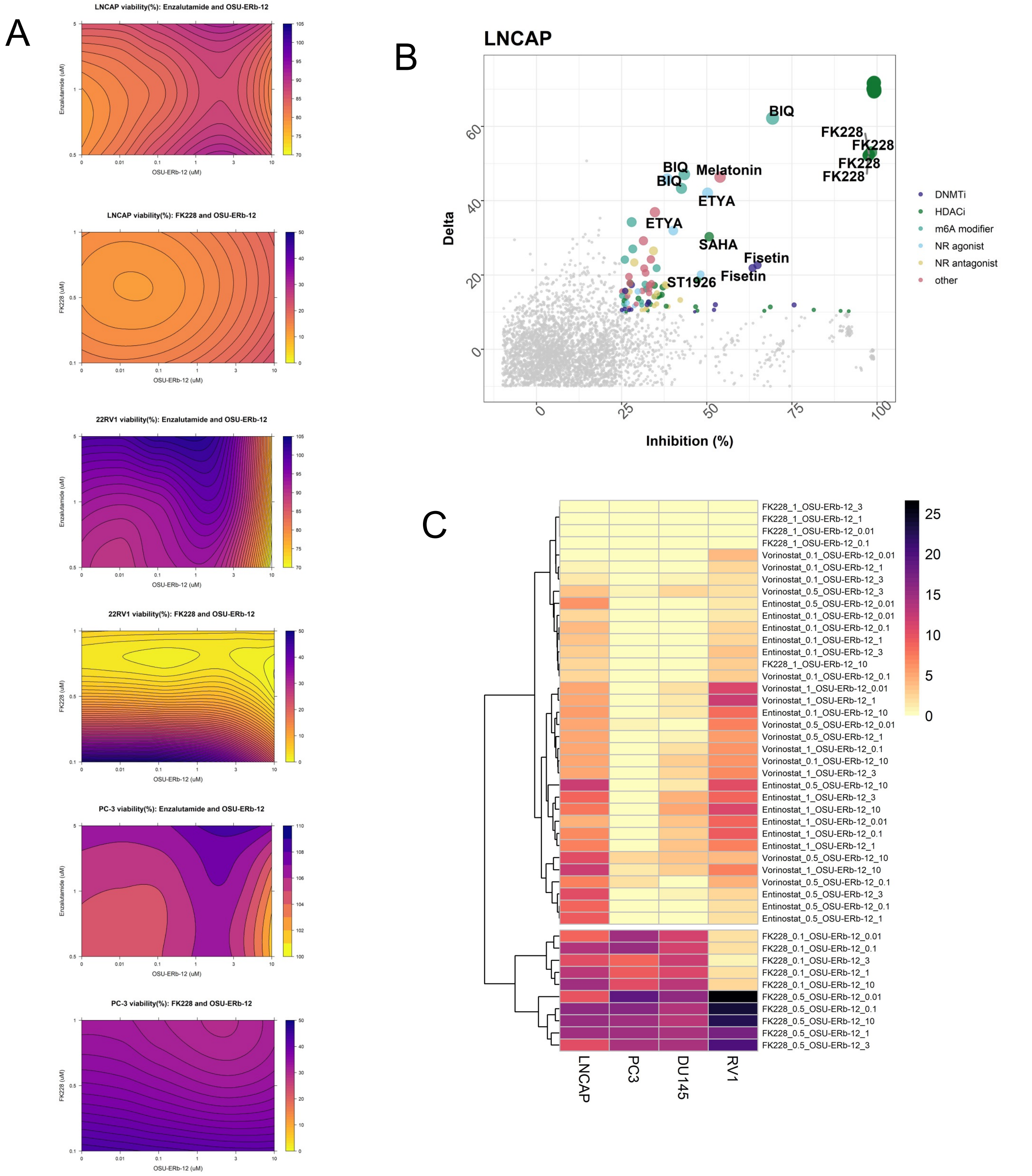

Supplementary Figure 2

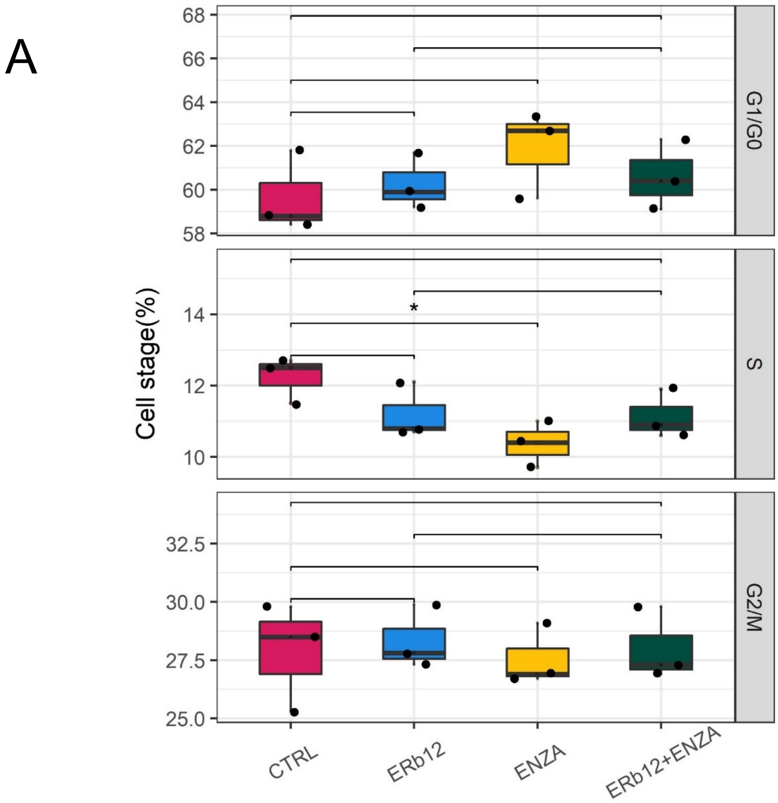

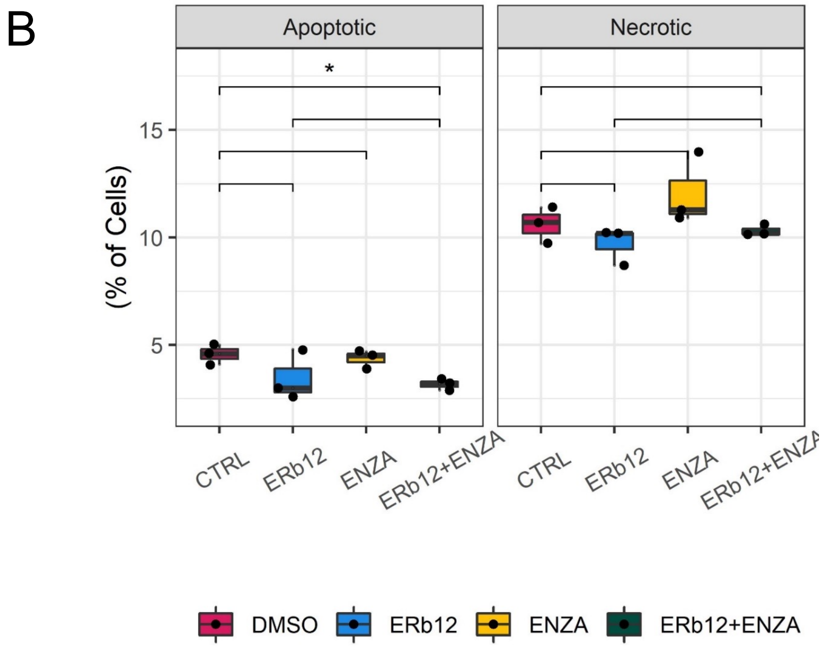

Supplementary Figure 3

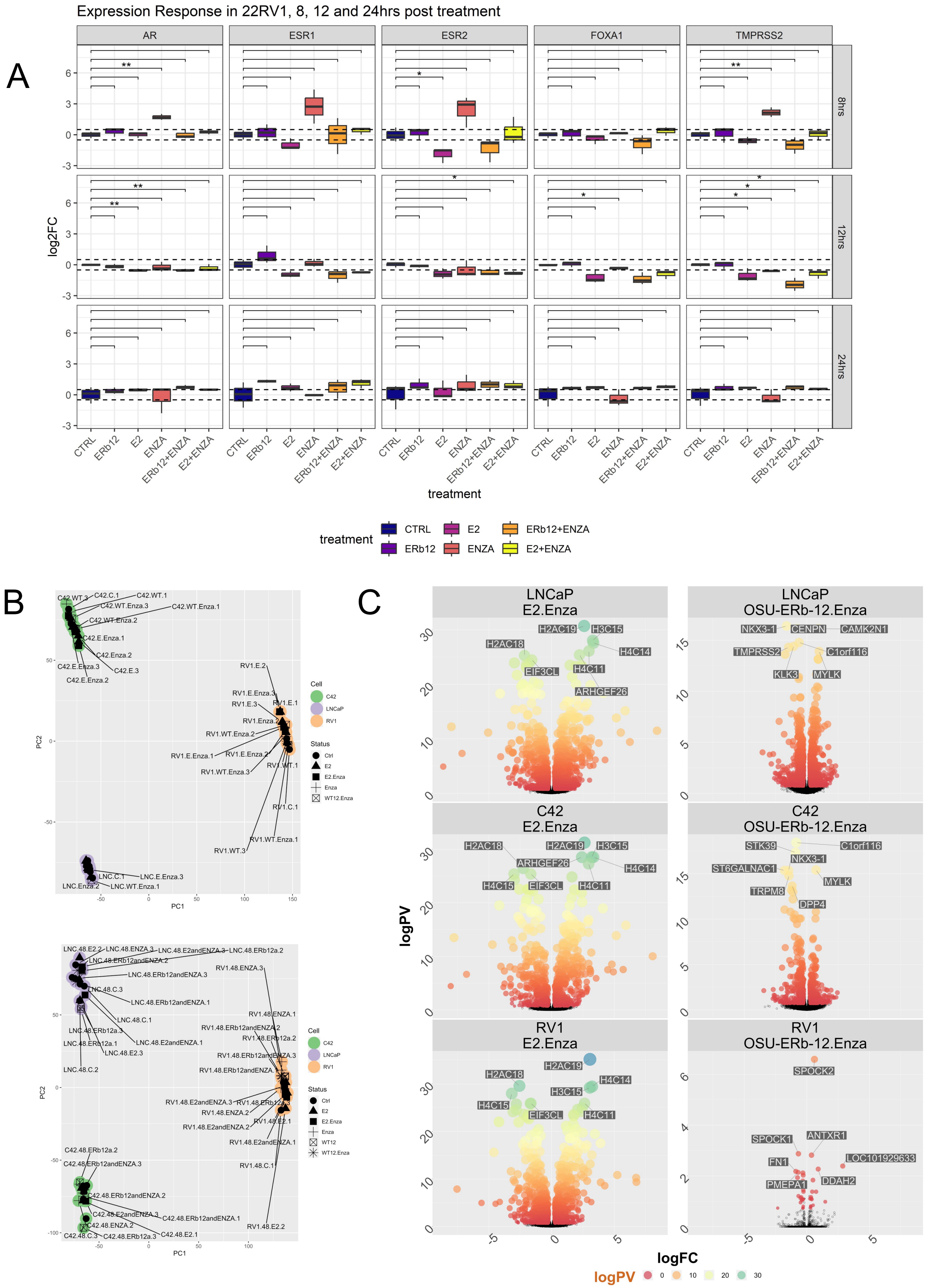

Supplementary Figure 4

A

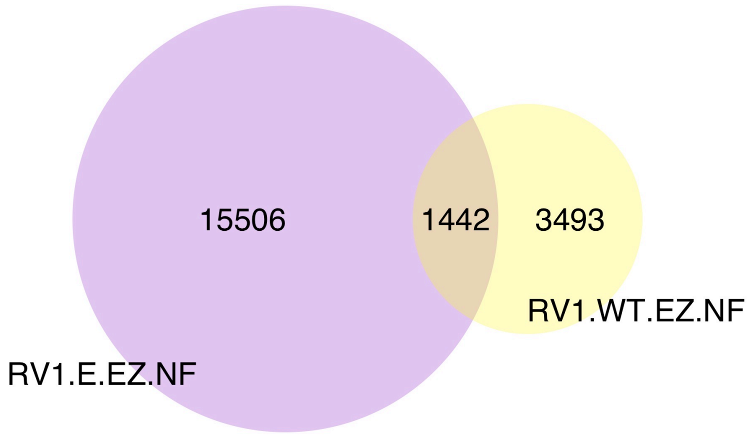

B

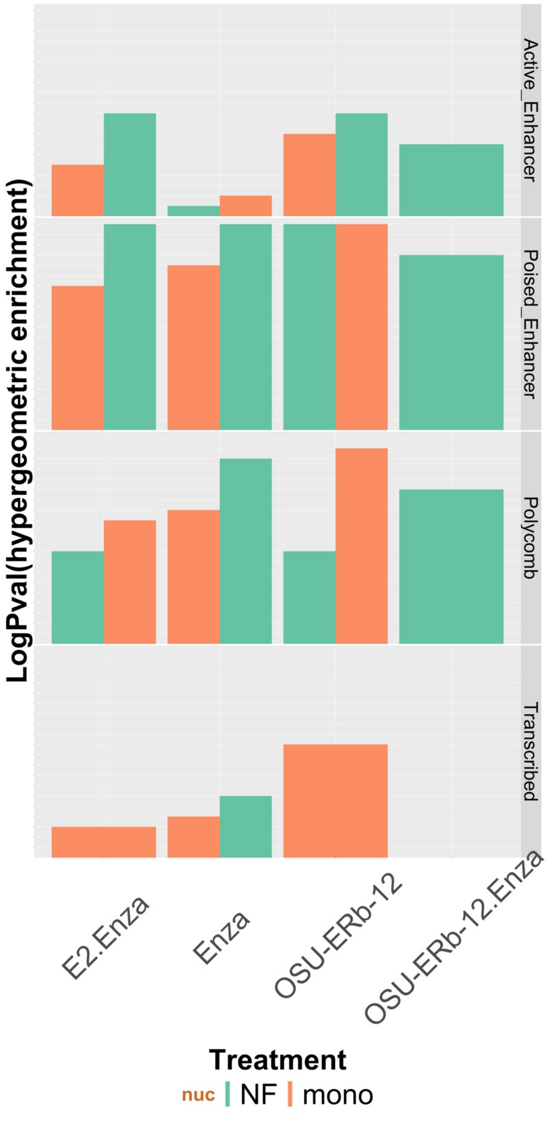

Supplementary Figure 5

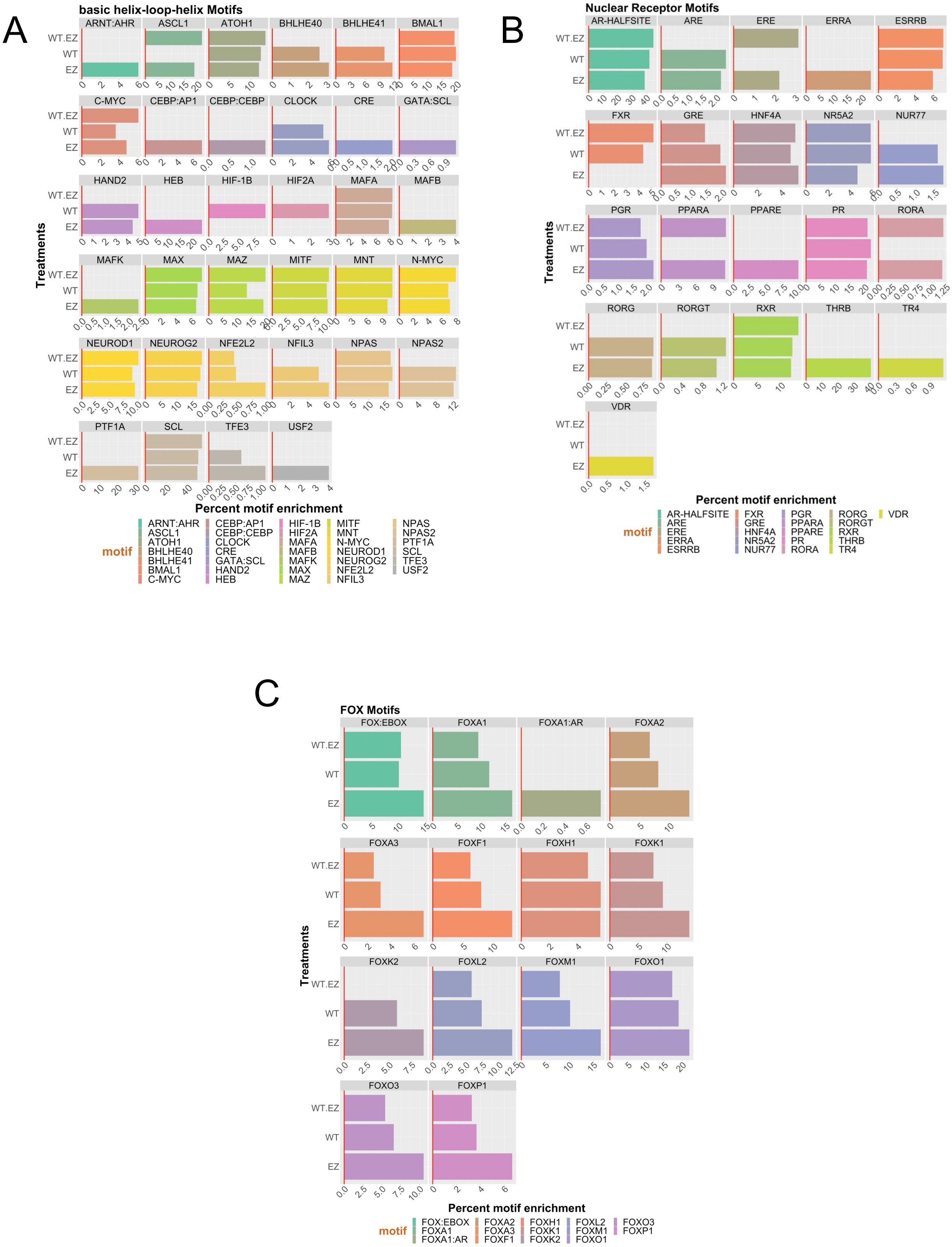

Supplementary Figure 6

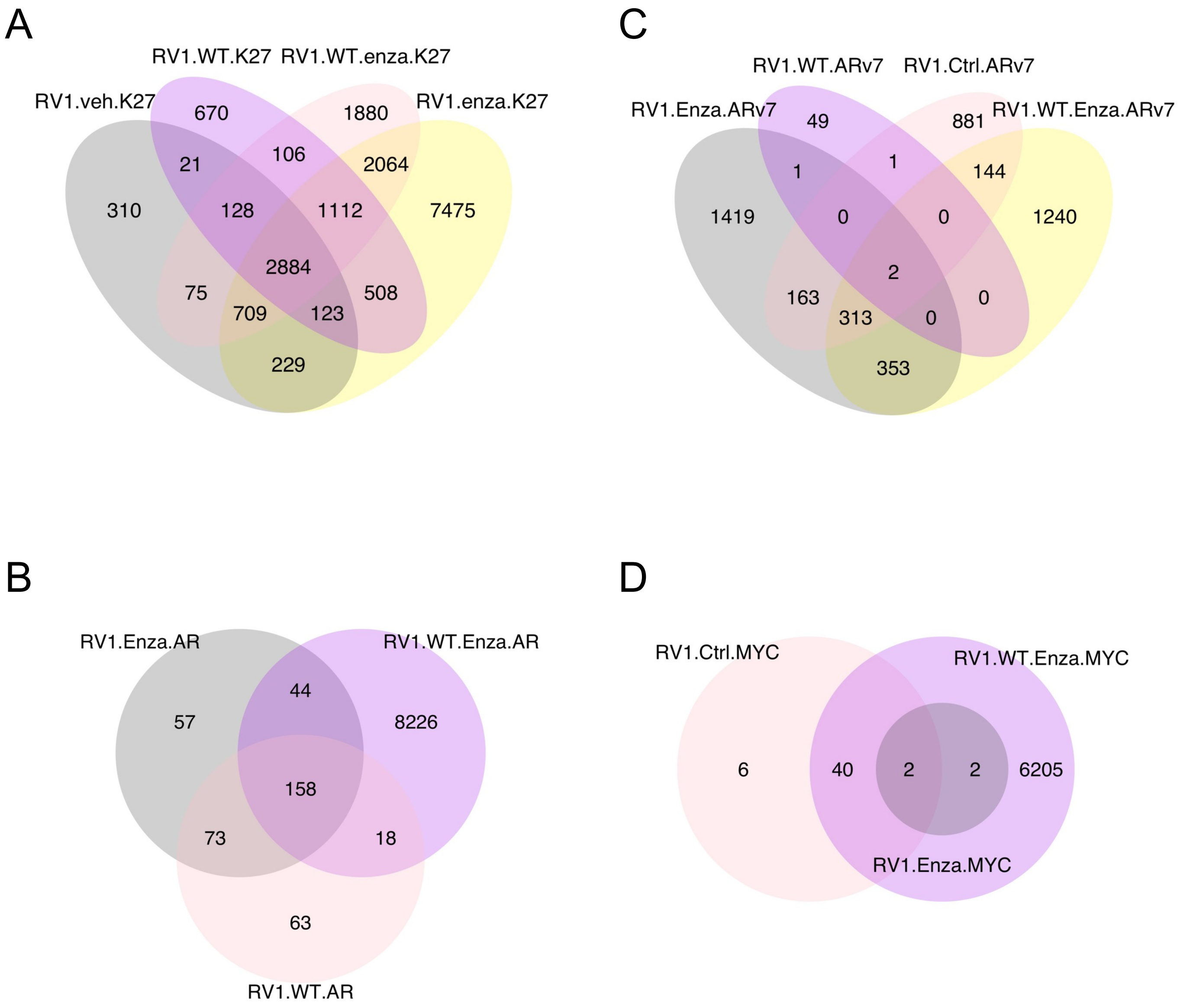

Supplementary Figure 7

A

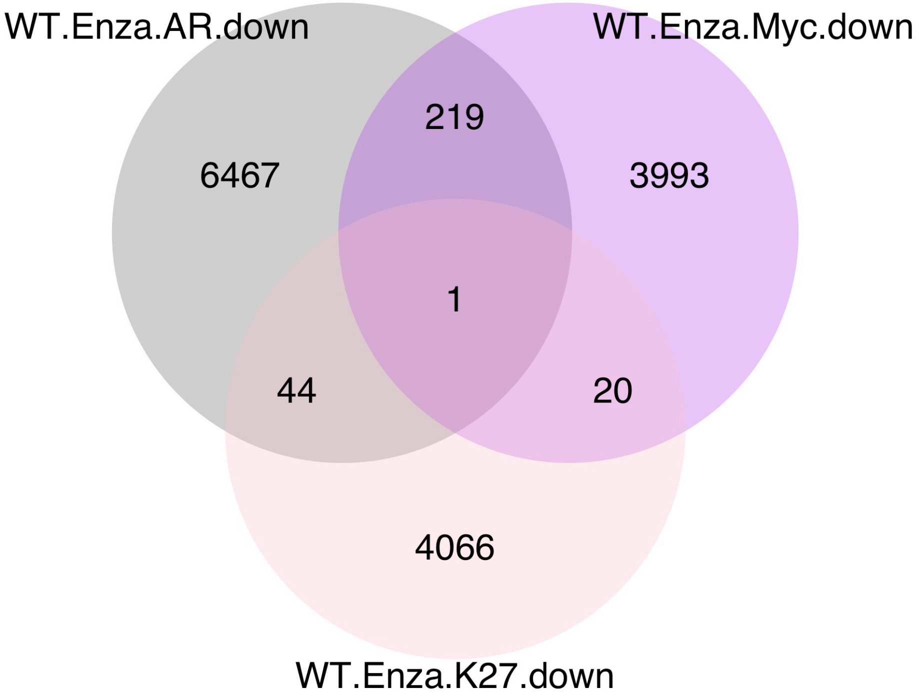

Supplementary Figure 8
